# The molecular landscape of cellular metal ion biology

**DOI:** 10.1101/2024.02.29.582718

**Authors:** Simran Kaur Aulakh, Oliver Lemke, Lukasz Szyrwiel, Stephan Kamrad, Yu Chen, Johannes Hartl, Michael Muelleder, Jens Nielsen, Markus Ralser

**Affiliations:** Molecular Biology of Metabolism Laboratory, The Francis Crick Institute, _1 Midland Road_, _NW1 1AT_, London, United Kingdom; Wellcome Centre for Human Genetics, Nuffield Department of Medicine, University of Oxford, United Kingdom; Department of Biochemistry, Charité – Universitätsmedizin Berlin, _Charitéplatz 1_, _10117_, Berlin, Germany; Berlin Institute of Health (BIH) at Charité – Universitätsmedizin Berlin, Berlin, Germany; Department of Biology and Biological Engineering, Chalmers University of Technology, _SE41128_ Gothenburg, Sweden; Shenzhen Institute of Synthetic Biology, Shenzhen Institute of Advanced Technology, Chinese Academy of Sciences, Shenzhen 518055, China; Max Planck Institute for Molecular Genetics, _Ihnestrasse 73_, _14195_ Berlin, Germany

## Abstract

Metal ions play crucial roles in cells, yet the broader impact of metal availability on biological networks remains underexplored. We generated genome-wide resources, systematically quantifying yeast cell growth, metallomic, proteomic, and genetic responses upon varying each of its essential metal ions (Ca, Cu, Fe, K, Mg, Mn, Mo, Na, Zn), over several orders of magnitude. We find that metal ions deeply impact cellular networks, with 57.6% of the proteome, including most signalling pathways, responding. While the biological response to each metal is distinct, our data reveals common properties of metal responsiveness, such as concentration interdependencies and metal homeostasis. We describe a compendium of metal-dependent cellular processes and reveal that several understudied genes can be functionally annotated based on their metal responses. Furthermore, we report that metalloenzymes occupy central nodes in the metabolic network and are more likely to be encoded by isozymes, resulting in system-wide responsiveness to metal availability.

## Introduction

Metal ions are integral to the functioning of biological systems with critical roles in biochemical reactions ^1^, vital metabolic pathways ^2^, protein evolution ^3^, and diseases like neurodegeneration ^4^, cancer ^5^ and microbial infections ^6^. As catalysts in enzyme active sites, reactant co-factors in redox reactions and by mediating protein-protein and protein-small molecule interactions, metal ions are required for energy transformation, biosynthesis, stress response and cellular signalling within prokaryotic, archaean and eukaryotic metabolic networks ^7^. Consequently, they are important for a wide range of biological processes, such as cell growth, protein folding, DNA repair, neurotransmission, and immune function ^8^.

The role of metal ions in the proteome is extensive. Budding yeast, arguably the best studied eukaryotic model system when it comes to metal ion biology, expresses at least 70 metal ion transporters and more than 800 of its ∼6000 confirmed proteins are annotated as metal-binders ^9^. Because the concentration of metal ions in the cellular environments is subject to constant fluctuations, cells sense, control, and buffer cellular metal ion concentrations against environmental fluctuations ^10,11^. However, the metal ion concentrations provided in the growth media for cells and tissues, are seldomly varied in molecular biology experiments. For example, only 0.8% (119 out of 14484, **Supplementary Figure 1a**) genome-wide yeast screens compiled by ^12^ deviate from the metal ion concentrations present in the standard growth medium.

Furthermore, even within this small subset, most screens explored metal ion toxicity, but not non-toxic concentration changes, i.e., those within a physiologically relevant range. Moreover, none of the screens covered metal depletion in minimal media devoid of amino acid supplements - which would be required for assessing the role of metal ions in biosynthetic metabolic pathways, in which metal-containing enzymes play a key role as catalysts (**Supplementary Figure 1b**).

Recently, individual studies have altered concentrations of metal ions such as zinc and iron in the yeast growth medium and conducted transcriptome and proteome analyses. The obtained data provides evidence for a widespread cellular responsiveness to metal ion perturbation ^13–15,16,17^. On the molecular level, ‘omic’ datasets that are obtained upon varying a metal ion concentration in the media, can however be challenging to interpret, for two main reasons. First, due to the extensive cellular concentration-buffering of the metal ions, a change in the media concentrations does not directly translate into a similar change in its cellular levels ^18–20^. Second, metal ion transporters are promiscuous, and altering the concentration of one metal ion will inadvertently influence the cellular levels of other metal ions, resulting in a complex relationship between metal availability in cultivation media, the intracellular concentration, and the response detected at transcriptome or proteome. The cellular responses to perturbations in metal availability are a combination of the concentration-buffering capacities, and the impact of the imposed environmental fluctuations and the interlinked intracellular changes ^10,21–23^.

Here, we aimed to create a resource that addresses the gap in knowledge about the role of metal ion concentrations in cellular networks. We varied the concentration of each typical metallic media component of minimal *S. cerevisiae* cultivation media systematically and over several orders of magnitude, resulting in 91 different growth media. We measured growth responses to the altered metal availability and employed a series of -omics technologies to monitor the biological responses: metallomics to quantify metal buffering and correlations between cellular metal concentrations, proteomics to measure the molecular response of cells, and growth screen of a genome-wide deletion library for identifying genetic interactions using metal depletion media. In parallel, we incorporated a dataset that captures the proteomic response to the deletion of each non-essential metal-related protein ^24^, and two metallomic datasets that quantify metal quantities in all strains from the *S. cerevisiae* knockout deletion and overexpression mutant collections ^25–27^.

Our comprehensive resource of metal biology unveils an extensive impact of metal ion concentrations across cellular biochemical networks and their regulatory landscape. We report that cellular metal responsiveness extends far beyond biomolecules and processes linked to metals due to direct metal-binding, metal-transport or metal-homeostasis activity. Instead, metal ion responsiveness affects a wide array of cellular processes, including transcription factors, signalling pathways, protein complexes and metabolic pathways that have not previously been implicated in metal ion biology. For example, we report that 28 out of 34 KEGG signalling pathways, including mTOR, contain metal-responsive proteins. While cells exhibit specific responses to the availability of each metal ion, we identify universal features of cellular metal responsiveness, such as cellular buffering capacities and the interplay between different metal ion concentrations. Moreover, we show that several hitherto uncharacterized genes induce characteristic metal-ion responses, and that these can be exploited to annotate their function. Lastly, we demonstrate how the high connectivity of enzymes and metabolites combined with the central location of metalloproteins within the metabolic network renders cellular metabolism remarkably sensitive to fluctuations in metal ions.

## 2. Results

We created nine series of cultivation media, in which the concentration of each metal salt typically supplied in synthetic minimal media, namely Calcium (Ca), Copper (Cu), Iron (Fe), Potassium (K), Magnesium (Mg), Manganese (Mn), Molybdenum (Mo), Sodium (Na), and Zinc (Zn), were varied one at a time, in 12-steps, over five orders of magnitude (**Figure 1a, Supplementary Note 1**). We chose to omit amino acid supplements in all our media formulations ^28^ to guarantee the activation of the cellular biosynthetic pathways, many of which require metal binding proteins ^29,30^. We then cultivated a prototrophic, haploid *S. cerevisiae* strain derivative of BY4741 ^31,32^ in each of the 91 media and the control in triplicates. To maintain consistency with previous literature and datasets, we refer to concentrations exceeding the typical media formulations ^33,34^ as “excess”, while those below as “depletion” (see **Supplementary Table 1** and **Methods**). Of note, we did not use chelators to completely deplete metals below trace concentrations present even in ultrapure laboratory solvents and materials, in order to avoid confounding chelator off-target effects ^35–37^ (**Supplementary Note 2**). To enable a precise determination of the amount of metal available to cells in each cultivation condition, we instead quantified the metal content in each of the 91 media using Inductively Coupled Plasma Mass Spectrometry (ICP-MS) (**Supplementary Figure 1c**, **Supplementary Table 2**).

**Figure 1.**
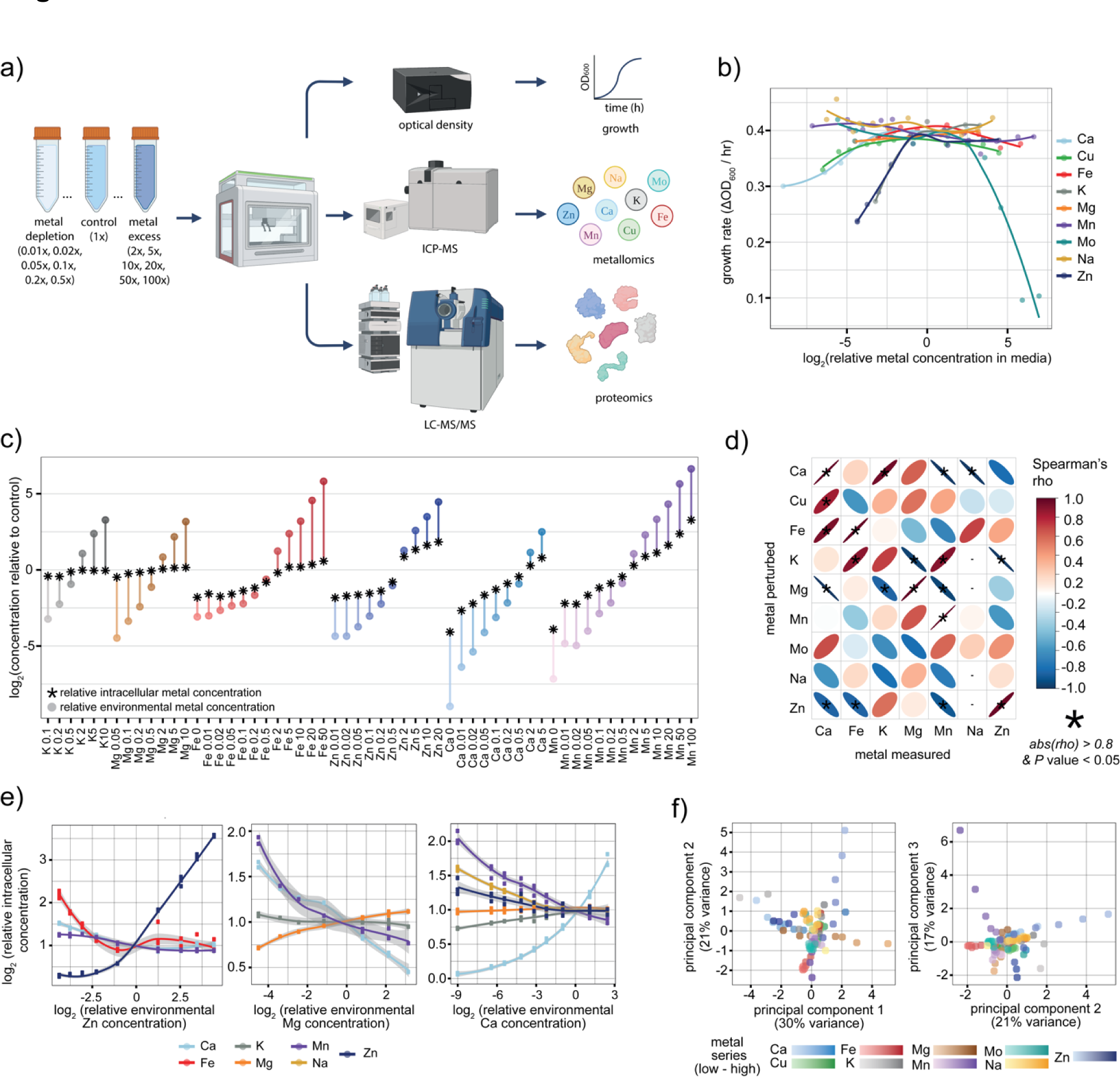
Metal ion homeostasis and concentration-interdependence a) Schematic: Experimental design for media generation, growth, metallomic, and proteomic characterisation of *S. cerevisiae* cultivated in 91 media, that constitute concentration gradients in its typical media supplements Ca, Cu, Fe, K, Mg, Mn, Mo, Na, and Zn. b) Growth rate of *a* prototrophic BY4741- derivative along each metal perturbation series. X- axis: log_2_(concentration of metal in media relative to control (synthetic minimal media)). Y-axis: growth rate (change in OD_600_ per hour, relative to time of inoculation) c) Quantification of the buffering capacity of cellular versus extracellular metal concentrations. X-axis: media concentration, relative to concentration in typical synthetic minimal media (control). Y-axis, coloured circles : log_2_(concentration of perturbed metal in cultivation media relative to that in synthetic minimal media (control). Y-axis, black stars : log_2_(cellular metal concentration of cells cultivated in each metal perturbation condition relative to that of cells cultivated in synthetic minimal media (control). d) Correlation between the metal concentration in cultivation media, and cellular concentration of other metals. Y-axis: metal that was perturbed in the cultivation media. X-axis: metal measured in *S. cerevisiae* cells cultivated in media with perturbed concentration of each metal. Colour indicates value of Spearman’s rank correlation coefficient. Black stars represent correlations that had a *P* value < 0.05 (calculated by cor.test() function in R). e) Examples of correlation between the environmental concentration of one metal (X-axis: log_2_(relative environmental metal concentration)) and the cellular concentration of another metal (Y-axis: log_2_(relative total cellular metal concentration)). f) Principal component analysis based on all measured total cellular metal concentrations separates a subset of samples according to the metal perturbed in the environment. Left panel: principal component (PC) 1 on the X-axis and PC 2 on the Y-axis. Right panel: PC 2 on the X-axis and PC 3 on the Y-axis. Colours indicate each unique cultivation condition with darker colours indicating high amounts of each metal and lighter colours representing low amounts.

*S. cerevisiae* cells maintained consistent growth across a high range of environmental (media) metal concentrations, in most concentration series (**Figure 1b, Supplementary Figure 1d**). However, growth rates reduced upon a ∼2-fold depletion of abundant metals K, Mg and Zn and at ∼8-fold depletion for Ca and Cu (**Figure 1b**). Despite successful Mo and Mn depletion to ∼8ppb (0.04 times the typical media concentration) and ∼2.8ppb (0.007 times the media concentration) respectively, no growth defects were observed, suggesting that these elements are either nonessential under the tested conditions, or required at extremely low amounts. Even though we omitted metal salts completely to prepare the lowest Fe and Na depletion media and used solvents and materials of the highest purity available, the Fe and Na concentrations in these media (8 ug/L of FeCl_3_ or 143 ppb of Fe atoms and ∼508 ug/L or ∼22100 ppb of Na atoms) were sufficient to sustain cell growth. At the other end of the concentration series, we observed that concentration changes of more than 5-fold (1 mg/L Na_2_MoO_4_) of Mo, more than 20-fold of Cu and Fe (0.8 mg/L CuSO_4_ and 5 mg/L FeCl_3_) resulted in a slowing of cell growth, indicating toxicity.

### Homeostasis is metal specific and involves concentration interactions

The observation that high magnitudes of environmental metal concentration changes are required to influence yeast cell growth is consistent with an extensive homeostatic machinery that allows cells to buffer cellular concentration against environmental changes ^10,21–23^. In order to generate a systematic dataset that captures the interdependency of cellular metal ion concentrations in relation to their extracellular levels, we quantified the total cellular concentration of Ca, Fe, K, Na, Zn and Mg (see **Supplementary Table 3**) using an ICP-MS protocol adapted from ^26^ . For WT cells cultivated in the standard condition the relative concentration of metals we observe are consistent with previous reports (Methods, **Supplementary Figure 1f**). The obtained concentration values reveal the ‘buffering range, the extracellular concentration span in which cells maintain a similar cellular concentration. Cells revealed the strongest concentration buffering for K, Mg, followed by Zn, Ca, and Mn (**Figure 1c**). Quantitatively, cells buffered at least 38-fold (Fe), 10-fold (K), 10-fold(Mn), 8-fold (Mg), 6-fold(Zn) and 3-fold(Ca) against excess, and 38-fold(Ca), 16-fold(Mg), 10-fold(Mn), 7-fold (K), 6-fold (Zn) and 2.5-fold(Fe) against depletion (**Figure 1c**, **Supplementary Table 4**).

Metal ions possess similar physical and chemical properties that result in promiscuity of metal transport systems ^10,11,38^. Thus, we next determined the relationship between the environmental concentration of one metal and the cellular concentration of any other metal. Seven out of the nine environmental metal concentration series affected the cellular concentrations of at least one other metal (based on Spearman’s rank correlation abs(rho) > 0.8 and p-value < 0.05, **Figure 1d, Supplementary Table 5**). Overall, cellular Mn and Ca levels were most sensitive to environmental concentration changes in other metals, while environmental K exerted the most widespread impact. For instance, changes in environmental K caused positively correlated concentration changes in cellular Fe (Spearman’s r = 0.88) and Mn (Spearman’s r = 0.89) and correlated negatively with cellular Mg (Spearman’s r= -0.96) and Zn (Spearman’s r ^=^ -0.98) levels. Perturbations in environmental Ca, Mg and Zn influenced the concentrations of three other metals each (**Figure 1e**), with cellular Mn concentration being negatively correlated with each of these three metals (Spearman’s r of -0.98, -0.94 and -0.88 respectively). Conversely, environmental concentrations of Cu and Fe positively correlated with cellular Ca concentration (Spearman’s r of 0.84 and 0.93 respectively). At least some of the observed interdependencies correspond to shared physical properties of metals and biochemical interactions; for instance, Mg-induced decreases in cellular Ca and Zn concentrations align with the promiscuity of divalent cation transporters ^39^. Despite these interactions, the metallomics profiles of samples from each cultivation media were sufficiently specific to group most samples in accordance with the metal perturbation in a principal component analysis (PCA) (**Figure 1f**).

### The proteome responds globally to changes in metal availability

We next conducted a quantitative proteomics experiment to capture molecular responses in the cells grown in the 91 conditions with altered metal concentrations, in triplicates. We employed a high-throughput proteomic pipeline that combines cell cultivation in multi-well plates, semi-automated sample preparation, microflow liquid chromatography, data-independent mass spectrometry data acquisition, and data processing using DIA-NN ^24,40,41^. After extensive filtering and quality control of the raw proteomics data (Methods), we obtained precise quantitative data for 2330 unique proteins. Of these, 1433 were quantified in at least 85% of all samples (**Methods, Supplementary Table 6**). For 3841 proteins in the *S. cerevisiae* proteome, quantitative copy number data has been estimated with an orthogonal strategy ^42^ which allowed us to estimate the proteomic mass represented by the observed proteomic changes. The overlap of 1916 proteins between our dataset and proteins quantified in ^42^ led us to estimate that our dataset quantifies around 87.9% of the proteome mass represented by these 3841 proteins with 76.8% of the proteome mass (corresponding to 1190 proteins) being quantified with over 85% completeness across the dataset (**Methods**). The average replicate coefficient of variation (CoV), which reflects the sum of technical and biological noise, was ∼15.7% with an average of 1837 proteins quantified per sample for the perturbation samples and 1871 proteins quantified per control condition sample. The biological signal in the dataset (average CoV of proteins across test conditions) was ∼27.9%, considerably higher than the noise levels.

To identify and classify differentially expressed proteins along the metal concentration series, we determined whether the relationship between environmental metal concentration and protein abundance was best represented by a null, 1^st^, ^2nd^, or 3^rd^-degree polynomial linear model using p-values from pairwise F-tests between each type of model (**Figure 2a**, **Methods, Supplementary Figure 2a**, **Supplementary Table 7**, **Supplementary Table 8**). We defined a protein as differentially expressed, if the p-value of the model that best represents the modelled relationship was < 0.05 and if difference between the minimum and maximum protein quantity along the environmental metal concentration series was at least 1.5-fold (i.e., *abs(max(log_2_(fold difference vs control)) - min(log_2_(fold difference vs control)) > log_2_(1.5)*. We used the same methodology to identify proteins altered along measured cellular (as opposed to environmental) metal concentrations, using binned metal concentrations and protein abundance data across the entire dataset, to identify protein-metal interactions resulting from the interdependency between cellular metal concentrations.

**Figure 2.**
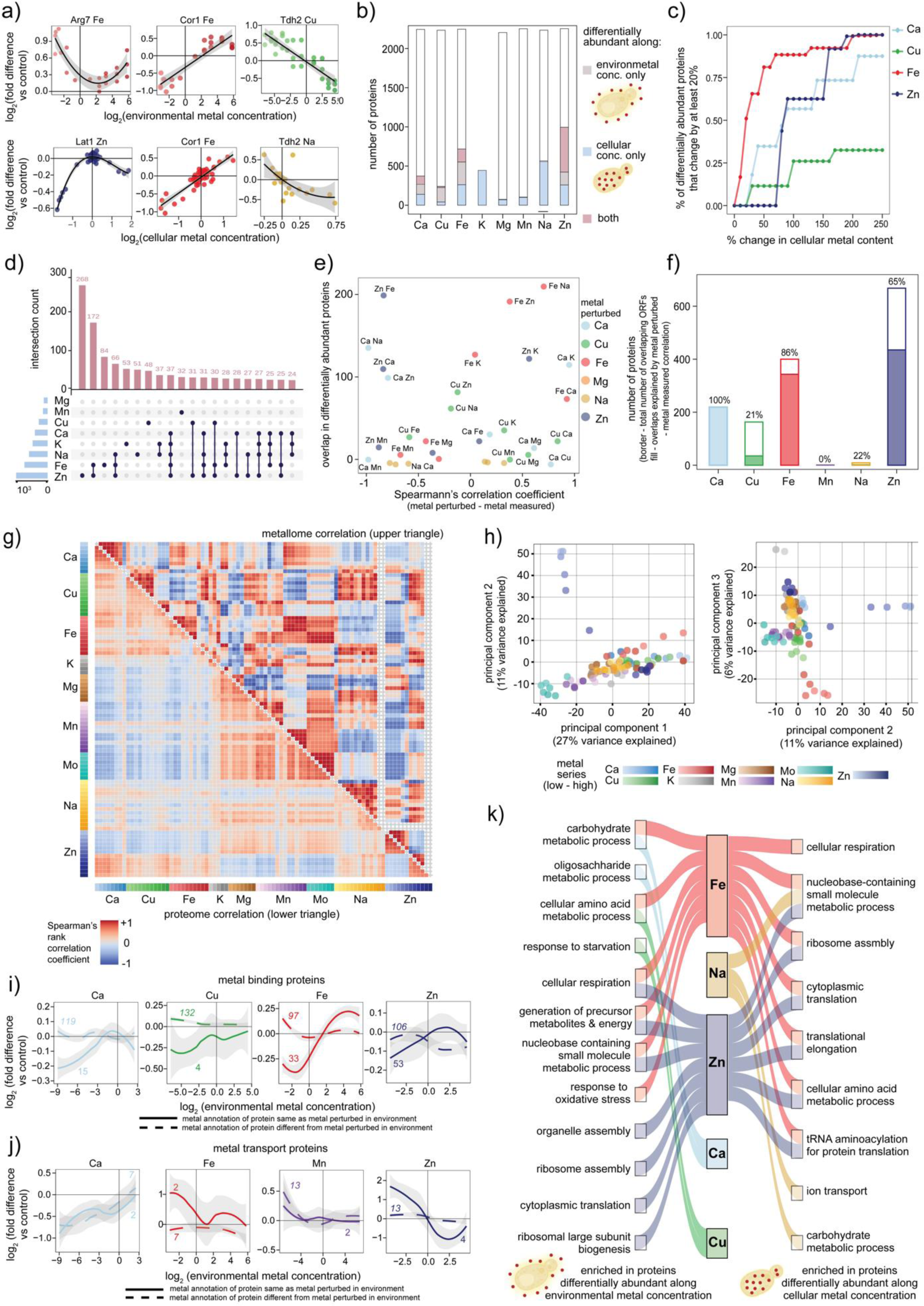
Identifying the metal responsive proteome a) Proteins that respond to metal ion contractions (selected examples that represent different metal ion - protein abundance relationships along environmental and cellular metal concentration series). Differentially abundant proteins were determined by conducting F-tests between null (y∼c), first (y ∼ mx+c), second (y ∼ mx^2^ + c) and third (y∼mx^3^ +c) degree polynomial models. Top row: proteins identified as differentially abundant based on polynomial fits of protein abundance relative to synthetic minimal media(control) (Y-axis) vs. log_2_(environmental metal concentration relative to control) (X-axis). The plots are labelled with the protein abundance shown, plus the varied metal ion concentration. Bottom row: proteins identified as differentially abundant based on polynomial fits of protein abundance relative to control (Y-axis) vs. log_2_(cellular metal concentration relative to control)(X-axis). b) Number of proteins identified as being differentially abundant along each metal concentration series (beige - along environmental metal concentration series, light blue - along measured cellular concentration and pink - along both environmental and cellular concentrations. c) Characterisation of thresholds of cellular metal concentration at which the proteome responds to metal perturbation. X-axis: percentage change in cellular metal content with intervals of 5%. Y-axis: percentage of total differentially abundant proteins along metal perturbation series that exhibit a change of at least 20% in abundance at each 5% increase (depicted by x-axis) in cellular metal concentration. d) Overlap between proteins differentially abundant along each metal perturbation series (either environmental or cellular). Pink bars in the upset plot indicate the size of overlap between conditions listed on the left. Light blue bars on the left indicate the number of proteins differentially abundant along each metal perturbation series. e) Relationship between number of differentially abundant proteins that overlap between environmental concentration series of each metal and cellular concentration of every other metal. X-axis: Spearman’s correlation coefficient calculated between environmental concentration of metal being perturbed and cellular concentration of each other metal. Y-axis: number of differentially abundant proteins that overlap between environmental concentration of each metal and cellular concentration of every other metal. The first metal in the text next to each point indicates the environmental (media) concentration series and the second indicates cellular (measured) concentration series. ‘*’ represents pairs that have a significant correlation between the metal perturbed and the metal measured based on metallomics data. f) Most overlaps between proteins differentially abundant along environmental concentration of one metal and cellular concentration of another are explained by perturbation of cellular metal concentrations. X-axis: metal perturbed in the environment (cultivation media). Y-axis: outer rectangle of bar plot indicates the total number of differentially abundant proteins along each environmental concentration series that overlap with differentially abundant proteins along any other cellular concentration series. Inner rectangle(fill) indicates the number of proteins that are differentially abundant along cellular concentration of another metal for which the correlation between the perturbed metal and measured cellular concentration is high (Spearman’s rho > 0.8 and P value < 0.05). g) Comparison of correlation coefficients between metallomics (upper triangle) and proteomics (lower triangle) profile of each pair of unique cultivation conditions. Colour indicates Spearman’s correlation coefficient with red indicating positive values and blue indicating negative values. h) Separation of each unique cultivation condition based on principal component analysis of the proteomics data along the first three principal components. Left panel: principal component (PC) 1 on the X-axis and PC 2 on the Y-axis. Right panel: PC 2 on the X-axis and PC 3 on the Y-axis. Colours indicate each unique cultivation condition with darker colours indicating high amounts of each metal and lighter colours representing low amounts. i) Average protein abundance of metal binding proteins in *S. cerevisiae* cells cultivated in media with a series of metal concentration variations. X-axis: log_2_(environmental (cultivation media) metal concentration, relative to that in synthetic minimal media control). Y-axis: log_2_(fold difference in abundance of protein relative to that in control). Solid line indicates proteins that are annotated in the Gene Ontology database to bind to the same metal as being perturbed in the cultivation media (indicated above each panel), dashed lines indicate proteins that are annotated to bind a different metal than that being perturbed in cultivation media. Numbers in normal font indicate the total number of proteins bearing annotations for the same metal being perturbed, those in italics indicate the total number of proteins that have an annotation that is different from the metal being perturbed. j) Average protein abundance of metal transporters in *S. cerevisiae* cells cultivated in media with a series of metal concentration variations. X-axis: log_2_(environmental (cultivation media) metal concentration, relative to that in synthetic minimal media control). Y-axis: log_2_(fold difference in abundance of protein relative to that in control). Solid line indicates proteins that are annotated in the Gene Ontology database to transport the same metal as being perturbed in the cultivation media (indicated above each panel), dashed lines indicate proteins that are annotated to transport a different metal than that being perturbed in cultivation media. Numbers in normal font indicate the total number of proteins bearing annotations for the same metal being perturbed, those in italics indicate the total number of proteins that have an annotation that is different from the metal being perturbed. k) Summary of the Gene Ontology - Slim (biological process) terms enriched in proteins differentially abundant proteins depending on the extracellular (left) and cellular (right) metal ion concentration. Rectangles in the central panel indicate the metal being perturbed in each series. Rectangles and annotations on the left indicate gene set terms that are enriched in proteins that were deemed as differentially abundant based on linear models fit to protein abundance along environmental (media) metal concentration. Colours represent each perturbed metal.

This analysis identified 1545 unique proteins, corresponding to 66% of the number of measured proteins and 81% in terms of the quantified protein mass, to be metal responsive. On average, 205 proteins were altered per metal perturbation series, when considering environmental concentration, and 342 when considering cellular metal concentrations (**Supplementary Table 6**, **Figure 2a**).). Overall, the most pronounced response was to Zn, with 164 proteins responding alongside changes in environmental Zn levels, 259 alongside cellular Zn levels, and 572 along changes in both environmental and cellular Zn levels (**Figure 2b**). On the other end of the spectrum, upon excluding the toxic concentrations at which we could not generate sufficient biomass for quantitative proteomics analysis, no proteins were found to be Mo-responsive.

Identifying K and Mg responsive proteins was technically challenging, as for these two metals only a limited environmental concentration range was testable (Supplementary Note 1). However, by using the cellular metal concentration changes, we identified 445 proteins for K and 71 proteins for Mg whose abundance correlated with the indirectly induced changes in cellular K and Mg, respectively. Similarly, despite only 12 proteins being identified as differentially abundant along the environmental Na concentration series, 555 correlated with indirectly induced changes in its cellular abundance (**Figure 2b**).

Across all metals, 1055 proteins were responsive to at least one environmental metal concentration change, and 1386 proteins were responsive to at least one cellular concentration change. Only 159 proteins were differentially regulated along at least one environmental concentration series only. For example, Arg7 protein abundance correlated with extracellular Fe concentration (**Figure 2a, top panel**). More proteins, 490, responded along cellular concentration changes only. For example, Lat1 levels changed according to the cellular Zn levels (**Figure 2a, bottom panel**). 712 proteins responded along both the environmental and the cellular series. For example, Cor1 responded to both changes in environmental and cellular Fe, while Tdh2 correlated with changes in environmental Cu and cellular Na (**Figure 2a**).

Next, we used our data to determine the critical concentration levels for initiation of a proteomic response, i.e., the minimum concentration changes at which cellular responses begin to be induced. We defined a threshold for protein level responsiveness as a change in protein abundance greater than 50% (*abs(log_2_(fold difference vs control)) > log_2_(1.5)*) and then evaluated the cumulative fraction of responsive proteins at every incremental 5% increase in magnitude of the measured cellular metal concentration (**Figure 2c**). This analysis revealed significant differences between the metal ions. For instance, a 75% alteration in the cellular Zn level was required to induce the proteomic response, while for iron, a 10% change in its cellular concentration was sufficient to induce the proteomic response (Figure **2c**).

### The proteome reflects concentration-interactions between metal ions

Previous studies have examined individual metal depletions of iron or zinc at the transcriptome ^13–15^ and proteome levels ^16,17^. While our data has a high correlation with the overall profiles published previously (**Supplementary** Figures 2b-2f), our results **(Figure 1d)** also suggested that proteomic responses might be induced by promiscuous changes in co-varying metal ion concentrations. Examining our data to quantify this property, we report that 761 proteins, or 49%, of all metal responsive proteins, showed protein abundance changes potentially resulting from covarying concentrations. To assess how unique the proteomic changes observed along each metal perturbation series were, we first visualised the intersections between proteins differentially abundant along each metal (**Figure 2d**). Four of the ten largest intersections correspond to proteins that are differentially abundant in a combination of two or more metals, while six correspond to proteins that are differentially abundant only along individual metal perturbation series — Zn, Fe, K, Na, Cu, and Mn (**Figure 2d**). Among these, proteins varying along the Zn series, followed by those changing along both Zn and Fe, and Fe alone triggered the largest responses (**Figure 2d**). Next, to assess how well the profiles of environmental metal perturbation series are explained by cellular changes in each other interacting metal, we compared the number of proteins differentially abundant along each pair of environmental and cellular metal concentration series to the correlation coefficients obtained from measured environmental (cultivation media) and cellular metal concentrations of each pair (**Figure 2e**). Some intersections resembled the metal-metal interconnections unveiled by the metallomics data. For instance, ∼200 proteins differentially abundant along the Fe series (environmental and cellular considered together) were also identified as significantly altered in abundance along the Zn or Na series (**Figure 2e**, red circles in top right and blue circle in top left). This overlap coincides with a high correlation coefficient between environmental Fe concentration changes and cellular Na and Zn concentrations as well as the link between environmental Zn concentration and cellular Fe concentration (**Figure 2e, x-axis**). Overall, Ca, the most interlinked metal based on metallomics data (**Figure 1d**), also showed a strong linkage with other metals at the proteome level with all the proteins identified as differentially abundant along the environmental Ca series that were also identified along the cellular concentration of any of the other metals being explainable by a metal-metal connection discovered via the metallomics data (**Figure 2f**).

We then evaluated how closely the proteome of a sample is related to its metal content by measuring similarity between the proteome and metallome profiles across all samples. We computed correlation coefficients between each unique pair of test condition samples (91) based on the metallome and proteome respectively and then compared the correlation coefficients for each pair of samples. Despite the generally low correlation between metallome and proteome similarities (Pearson correlation coefficient = 0.24 and Spearman rho = 0.18), specific conditions, such as Mo excess and Mn excess, or Cu depletion and Zn excess, showed clear correlations (**Figure 2g**). Principal component analysis (PCA) of the proteomics dataset revealed distinct separations for Zn depletion, Fe depletion, and K depletion samples along the first 3 components alone, with the remaining samples being arranged in slightly overlapping, but biologically relevant clusters of conditions (**Figure 2h**). A PCA analysis of the entire proteomics dataset showed that unlike PCA results of the metallomics dataset, which separated the Mg depletion and Mn excess out along PC1 and PC3 respectively, the Mn excess proteomes were clustered near the Mo and control condition sample and the Mg depletion samples were placed centrally amongst samples from other conditions.

### Cells cultivated along metal gradients reveal a compendium of cellular responses

Metal binding proteins and metal transporters represent the most well studied proteins in relation to metal ion biology. Indeed, many of the proteins differentially expressed along the metal concentration series are metal binding proteins, and these frequently respond to changes in the availability of the metal they bind (**Figure 2i**). However, we observed substantial differences between the metal ions. While most Fe- and Ca binding proteins responded strongly to Fe and Ca depletion, respectively, Zn-binding proteins were less responsive to Zn depletion (**Figure 2i**). Although we could only quantify a small number of metal transporters, we observed a different metal specific behaviour in this category: Fe, Zn, and Mn transporters were specifically responsive to the depletion of their interacting metal, while the abundance of the two quantified Ca transporters decreased at high calcium levels, indicating the presence of a negative feedback response (**Figure 2j**). Intriguingly, the abundance of metal transporters annotated to specifically transport metals other than Ca also decreased in Ca depletion.

To explore cellular processes responsive to changes in metal availability at a broader scale, we conducted a gene-set enrichment analysis using the Gene Ontology (GO), GOslim and KEGG databases (Methods). Cellular respiration, translation and transcriptional processes, stress response pathways, metabolic pathways as well as ion homeostasis processes were overrepresented among proteins differentially abundant along metal concentration series (**Figure 2k)**. Our analysis recapitulated known metal-specific molecular functions. For example, Fe binding, Fe-S cluster binding, heme binding proteins as well as enzymes with oxidoreductase activity were enriched along the environmental Fe perturbation series, lyase activity and oxidoreductase activity along Cu series and many ribosomal and oxidoreductase processes along the Zn series (**Supplementary Figure 2g**, **Supplementary Table 9**).

The enrichment analysis also revealed the involvement of less well documented responses to changes in metal ion availability. For instance, we observed a broad crosstalk between metal ion concentrations and cellular signalling pathways. In 28 out of 34 signalling pathways (GO-biological process annotations) for which proteins were quantified in our dataset, at least one protein was responsive to a metal ion concentration (**Supplementary Table 10)**. These included known metal response pathways, such as the calcineurin signalling pathway ^43^, both quantified proteins of the osmosensory pathway ^44^, but also signalling pathways with other canonical functions. For instance, all four quantified proteins of the G protein-coupled receptor pathway ^45^ and four of the five proteins quantified that map to protein kinase A signalling ^46^ were differentially expressed in metal perturbation media. Notably, our dataset reveals a strong metal response in the abundance of the mechanistic target of rapamycin (mTOR) pathway, which to the best of our knowledge has not been associated with metal ion responsiveness thus far. Seven out of eight quantified mTOR related proteins (Sit4, Ksp1, Kog1, Slm1, Stm1, Tap42, Tip41) were differentially expressed upon changes in metal concentrations. For example, Kog1, a subunit of the TORC1 responded to Cu, Fe and Zn availability in cultivation media while Sit4 responded to cellular concentrations of Fe and Na.

Due to their low abundance, our dataset quantified only 22 transcription factors. Twelve of these, including Yap1 which has a known role in Fe homeostasis ^47,48^ and Zn-finger or Zinc cluster transcriptional activator such as Cat8, Gat1 and Gts1 were differentially expressed in at least one metal perturbation series.

We could quantify at least one protein from 340 known protein complexes (GO-cellular compartment annotation) and observed a metal ion response in 289 of these (**Supplementary Table 11**). For 145 complexes, we quantified at least 75% of the components (**Supplementary Table 11**). In 128 of these (∼88%), at least one component was metal-responsive with 112 (∼77%) showing a change in at least 50% of their components. All the 38 large (five or more proteins involved) complexes (including the proteasome, vacuolar proton-transporting V-type ATPase, retromer complex, mannan polymerase and GPI-anchor transamidase complex) that were quantified with over 75% coverage contained at least one protein that was affected by at least one metal perturbation.

Finally, our dataset revealed a particularly strong metal responsiveness within the metabolic network. In 38 of the 39 KEGG metabolic pathways, for which we quantified more than 75% of the enzymes, at least one enzyme was differentially expressed along the metal concentration series. In 35 of these pathways, at least 50% of the quantified enzymes were metal responsive (**Supplementary Table 12**). Highly metal responsive KEGG pathway terms include steroid biosynthesis (85% proteins responsive), glycolysis (75%), TCA cycle (72%) and biosynthesis of secondary metabolites (81%). All proteins mapping to fatty acid biosynthesis and elongation (5), histidine metabolism (12), thiamine metabolism (4) and propanoate metabolism (9), 14 out of 15 enzymes of tryptophan metabolism and 10 out of 11 enzymes of lysine biosynthesis pathway were differentially expressed alongside a metal concentration series. Notably, the only four KEGG terms which we quantified at a high coverage but for which we did not observe a high responsiveness, are indirect participants in the metabolic network (i.e., ABC transporters, protein export, RNA polymerase), thus, essentially, all captured primary metabolic processes were metal responsive.

### Metal responsiveness clusters proteins according to function

Previous work , including our own studies on the yeast metabolome (Mülleder et al. 2016) and proteome (Messner et al. 2023) have revealed that clustering of ‘omic’ profiles can be an efficient strategy for protein functional annotation. To identify groups of proteins that respond in a similar manner along the metal concentration series, we employed an ensemble clustering approach ^49^.

We clustered the proteomics data in two parallel pipelines. In the first, we clustered the proteomes of cells cultivated in each individual metal series separately, while in the second, we clustered all proteomics data obtained in the Ca, Cu, Fe, Mg, Mn, Mo, K, Na and Zn series together. For the former (metal-wise clustering), only proteins that were identified as differentially abundant along each metal series were retained for that specific metal, whereas for the latter (all- metal clustering), all proteins detected in at least 85% of the entire dataset were included. In both instances, we utilised three clustering algorithms — density-based CommonNN ^50,51^, spatial k-Means(++) ^52,53^, and a community-detection algorithm (Leiden ^54^). Then we integrated the co-clustering matrices into a singular matrix with equal weighting, followed by a final hierarchical (Ward’s method ^55^) clustering step to define the final clusters (**Supplementary Figure 3a**). We obtained a total of 96 fuzzy clusters (with a range from 4 clusters for Mg and Mn to 27 clusters along the Zn concentration series) from the metal-wise clustering pipeline and 35 clusters from all-metal clustering. The coarse structure of the clustering is mainly driven by the Leiden-, the fine structure by the CommonNN and the kMeans-clustering. Functional enrichment analysis using the Gene Ontology, GO Slim, KEGG and Enzyme Commission databases were conducted for each cluster (representative examples are shown in **Figures 3a & b**, and a summary in **Figure 3c)**. Twenty of the 35 all metal clusters and 26 of the 96 metal wise clusters, cumulatively representing ∼60% (1061/1764) of the proteins, were enriched in at least one functional term (**Supplementary Table 13**). For example, a cluster of 29 proteins displayed an increase in abundance at low Cu concentrations (**Figure 3a**). These include enzymes of amino acid biosynthesis (Aat2p, Arg4p, Arg5,6p, Aro1p, Aro2p, Aro4p, Asn1p, Bat1p, His1p, His5p, Lys21p, Hom2p, Hom3p, Ilv1p, Ilv2p, Ilv3, Leu4, Lys1p, Lys2p, Lys21p Trp2p, Trp5p), the Lysyl-tRNA-synthetase Krs1p and others critical for mitochondrial function (Ggc1- the mitochondrial GTP/GDP transporter) and metabolism (Pyc2 - a pyruvate carboxylase that aids in the maintenance of precursors for the TCA cycle through the anaplerotic conversion of pyruvate to oxaloacetate in the cytoplasm), reflecting the key role of Cu in mitochondrial respiratory chain proteins that enable the production of amino acid precursors. Another cluster obtained via the all- metal clustering pipeline identified a group of 90 proteins characterised by a complex profile, involving protein abundant changes along the Ca, Zn, Mn, Cu and Fe series. The cluster was enriched for terms related to cation transport activity (**Figure 3b**).

**Figure 3.**
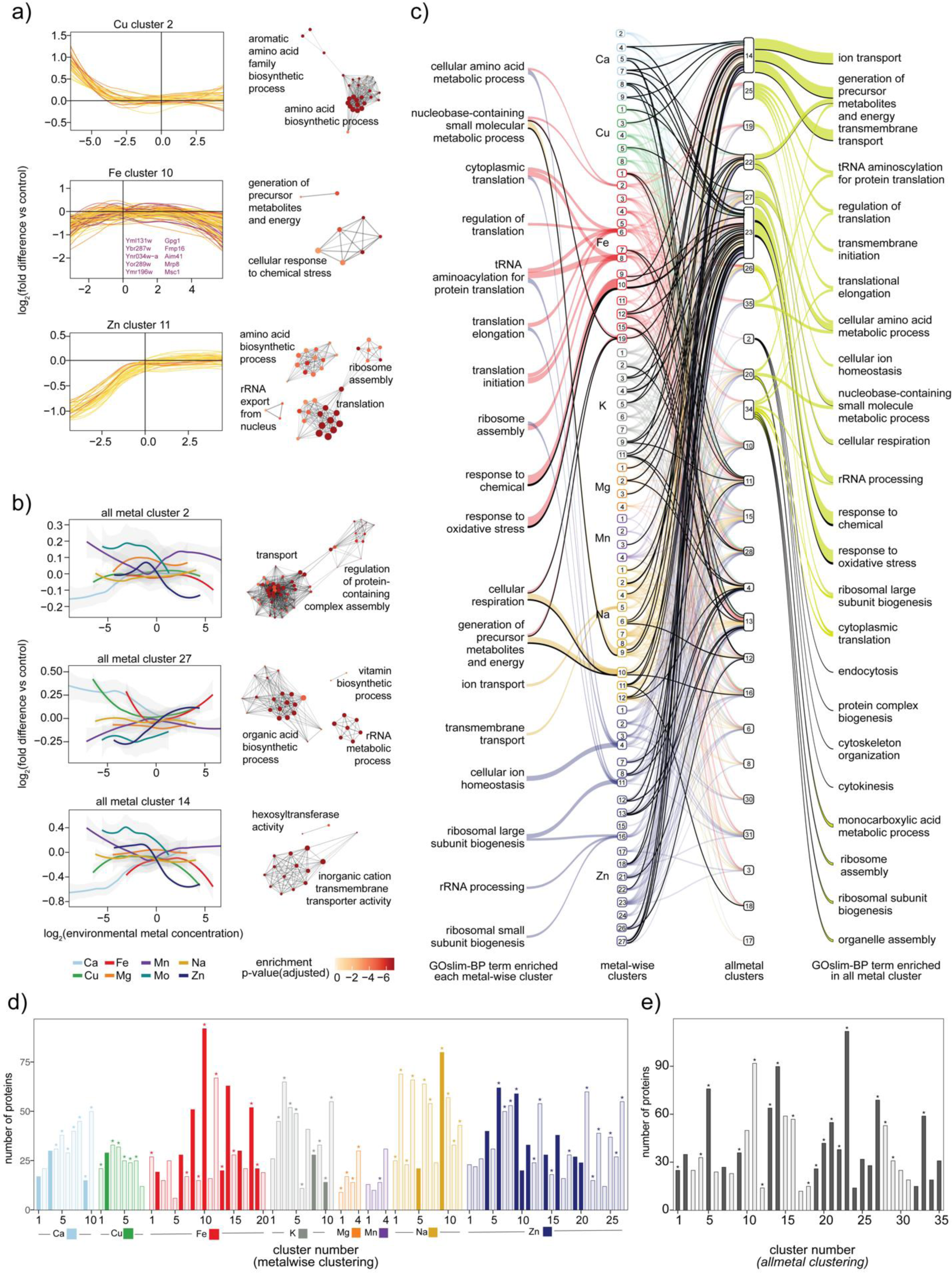
Metal responsiveness clusters proteins according to function a) Metalwise Ensemble-clustering. Examples of different clusters of proteins that show a similar response alongside a metal concentration gradient. Each panel depicts the log_2_ (environmental metal concentration relative to synthetic minimal media control) on the X-axis and log_2_(fold difference of protein abundance relative to that in control) on the Y-axis with colour indicating UniProt annotation score. The names of any protein bearing UniProt annotation score < 3 (poorly characterised proteins) are indicated in the figure. To the right of each graph in each horizontal panel is a network plot of all the Gene Ontology biological process terms that overrepresented the cluster. Each circle represents an individual geneset term with its colour representing the adjusted p-value of enrichment and size corresponding to the number of proteins mapping to the term. Gene sets with shared proteins are connected by grey lines. Groups of clusters are annotated with names that summarise groups of gene set nodes as determined and visualised using the *aPEAR* R library. b) Ensemble clustering across all metal ion gradients: Examples of different clusters of proteins that show a similar response alongside all metal concentration gradients. For each panel, X-axis: log_2_ (environmental metal concentration relative to synthetic minimal media control) and Y-axis: log_2_(fold difference of protein abundance relative to that in control) averaged for all proteins in the cluster for each metal perturbation series. Colour indicates the metal perturbation series and grey zone along the coloured lines indicate the 95% confidence interval around the mean abundance of all proteins in each metal perturbation series. To the right of each graph in each horizontal panel is a network plot of all the Gene Ontology biological process terms that overrepresented the cluster. Each circle represents an individual geneset term with its colour representing the adjusted p-value of enrichment and size corresponding to the number of proteins mapping to the term. Gene sets with shared proteins are connected by grey lines. Groups of clusters are annotated with names that summarise groups of gene set nodes as determined and visualised using the *aPEAR* R library. c) Summary of the Gene Ontology Slim (biological process) terms enriched in each cluster obtained via ensemble clustering. The two columns in the middle indicate cluster number after ensemble clustering - metalwise clusters are on the left and allmetal clusters on the right. The links between the clusters represent proteins with coloured links representing proteins belonging to each metalwise cluster that have a UniProt annotation score >2 and those that are black representing proteins with UniProt annotation score < 3. Gene set term names connected by links to the left of each metal wise cluster represent GOslim-BP terms enriched in each metalwise cluster while terms to the right of the allmetal clusters (connected by light yellow-green coloured links) represent GPslim-BP terms enriched in each allmetal cluster. Only those proteins that are part of a cluster (metalwise or allmetal) that has at least one GOslim-BP term enriched are included in the figure. d) Number of poorly characterised proteins in each cluster obtained via ensemble clustering (*metalwise-clustering* branch). X-axis indicates a cluster identifier assigned to each cluster that was obtained after ensemble clustering (using CommonNN, k-Means, and Leiden methods for each individual clustering step and hierarchical clustering on the co-clustering matrix to define the final clusters) of proteomics data from each metal perturbation series separately. Y-axis: number of proteins in each resultant cluster with the filled bars indicating whether any gene function term was enriched in the cluster and a star (*) above the bar indicating that at least one poorly characterised protein (UniProt annotation score < 3) is present in the cluster. e) Number of poorly characterised proteins in each cluster obtained via ensemble clustering (allmetal-clustering branch). X-axis indicates an arbitrary cluster number assigned to each cluster that was obtained after ensemble clustering (using CommonNN, k-Means, and Leiden methods for each individual clustering step and hierarchical clustering on the co-clustering matrix to define the final clusters) of the entire proteomics dataset (proteomes from all metal perturbation samples). Y-axis: number of proteins in each resultant cluster with the filled bars indicating whether any gene function term was enriched in the cluster and a star (*) above the bar indicating that at least one poorly characterised protein (UniProt annotation score < 3) is present in the cluster.

Overall, 23% (26/110) of the ORFs placed in clusters with enriched for GO-molecular function terms were related to metal binding functions, ∼49% (78/227) were placed in clusters enriched for mitochondria-related and ∼69% (157/227) in clusters enriched for ribosome related function. In addition, a range of GO-biological process and KEGG pathway terms not usually linked to metal ions, like the organisation of the cytoskeleton and the assembly of organelles (**Figure 3c, Supplementary Table 13**), were also enriched in numerous clusters.

### Incorporating functional genomics datasets to elucidate protein function

Notably, we found that 44 of 72 poorly characterised proteins (UniProt annotation score <3) captured by our proteomes were assigned to clusters enriched for a functional term (**Figure 3d & e**). To evaluate whether these associations provide relevant and reliable functional information for these proteins, we incorporated additional and complementary, genome-scale datasets relevant for metal biology. First, we cultivated a genome-wide yeast knock-out collection, consisting of 4850 single-gene deletion mutants in a prototrophic derivative of the genome-scale gene deletion mutant collection of *S. cerevisiae* ^29,56^ on 16 different metal omission media (depleted of Ca, Cu, Fe, Mn, Mo and Na, containing three concentrations of K, Mg and Zn, and one additional Fe depletion media that was generated using the metal chelator (2’-2’ bipyridyl ^57^) (**Methods**). We then measured colony sizes after 48 hours of growth using flatbed scanners and the *pyphe* toolbox ^58^. In total, we collected 357,972 colony size measurements (**Methods**) from which we calculated effect sizes and *P* values *((abs(mean effect size) > log(1.2)* and an adjusted *P* value *< 0.10* upon multiple testing correction using Benjamini-Hochberg)) for the growth of each mutant under each cultivation condition. This approach identified 734 genetic interactions with metal ion availability, involving 642 unique gene deletions, among the 4759 tested knockouts (**Figure 4a, Supplementary Table 14, Methods**).

**Figure 4.**
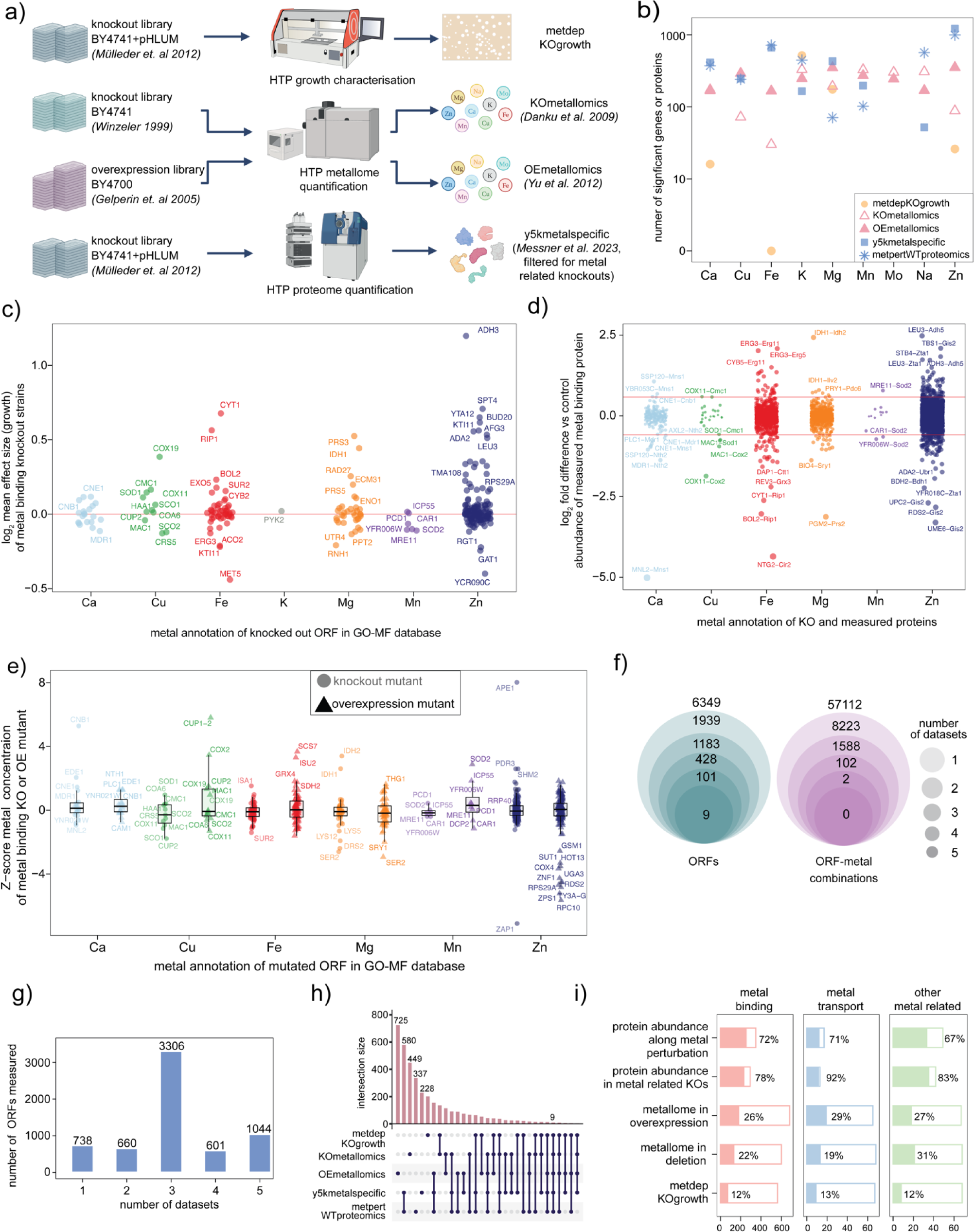
Data integration to generate a comprehensive resource for studying metal responsive proteins a) Data integration: Summary of a new genetic interaction dataset, two metallomic datasets of gene deletions and gene overexpression, respectively ^25,111^, and a genome-wide proteomic dataset ^24^ that integrated with the metallome and proteomic data, to create a comprehensive resource about metalloprotein function. b) *Number of Open Reading Frames (ORFs) identified as significantly affected across the five datasets* as summarised in a). X-axis : metal that was perturbed or in the case of the proteomic dataset of genome wide deletion mutants ^24^ - metal that was connected (based on GO-MF annotations) to the gene being deleted. Y-axis: number of genes or proteins that were identified as significantly affected. Colour indicates the type of assay (yellow – fitness inferred from end point colony size measurements, pink - cellular metal concentration, blue - proteomes). Shape indicates each individual dataset. c) *Effect of deletion of ORFs annotated for metal binding on the growth of S. cerevisiae on metal depletion agarose media*. X-axis: metal that was depleted and the metal-binding annotation of the ORF. Y-axis: log_2_(effect size (mean colony size of replicates of mutant relative to that of control, divided by the standard deviation of colony sizes of the mutant) - as determined by *pyphe-analyse*), averaged across all replicates. Colour indicates both the metal that was depleted and the metal binding annotation as the dataset was filtered to retain values where the metal depleted matched the metal binding annotation of the mutant. d) *Effect on protein abundance of metal-binding proteins upon deletion of ORFs encoding other proteins annotated to bind the same metal.* X-axis: the metal annotation of the deletion mutant as well as the measured protein. Y-axis: log_2_(fold difference of protein abundance in the deletion mutant vs. WT *S. cerevisiae*). Colours indicate the metal annotations of the deleted genes and measured proteins. Labels next to some of the points indicate the gene deleted (in capitals) followed by the protein measured (in title case). Only those points with a log_2_(fold difference vs control) > log_2_(1.5) are labelled using the geom_text_repel() which does not plot unavoidable overlaps. e) *Effect of deletion and overexpression of ORFs encoding metal-binding proteins on the cellular concentration of the metal each ORF is annotated to bind in the GO database*. X-axis: metal annotation of each ORF and metal quantified in each mutant. Y-axis : Z-score of metal concentration in each mutant (data from ^25,26^, Z score calculated by ^27^). Colour indicates the metal annotations as well as metal measured. Shape of points indicates the type of mutant - circles indicate knockout deletion mutants and triangles indicate overexpression mutants. f) *Intersection between ORFs (left) and ORF-metal combinations (right) that were identified as a significant hit across the five datasets.* Size of circle indicates how many datasets are considered with the largest circle representing the set of ORFs or ORF-metal pairs that are significant in any one dataset and the smallest representing those that are significant in all five. Numbers at the top, outside the largest circle represent the total number of unique ORFs and ORF-metal pairs that were measured in all five datasets, cumulatively. g) *Intersection between ORFs assayed across the five datasets*. X-axis: number of datasets an ORF was assayed / measured in. Y-axis: number of ORFs. h) *Intersections between ORFs that were a significant hit in any of the five datasets (Upset plot).* Black circles and lines between them indicate the identity of the datasets in the overlap, pink bars on top indicate the number of ORFs that are shared between the datasets indicated by the black circles and lines. i) *Metal related proteins identified as significantly affected in each dataset.* Panels represent the type of annotation a protein has in the Gene Ontology -Molecular Function database. X-axis: number of ORFs (outer bar outline represents the total number measured or assayed in each dataset and the inner filled-up bar represents the number that was significantly affected in at least one metal perturbation condition or metal-related mutant). Y-axis: dataset name and description.

The identified genetic interactions were enriched for metal protein binding, endosomes, protein complexes, ribosomes, translation, mitophagy, amino acid, amide and peptide biosynthetic pathways (**Supplementary Table 15**). At the individual metal level, a high number of genetic interactions were discovered for K (516) and Mg (175), followed by Zn (26) and Ca (16) (**Supplementary Figure 4a** and **Figure 4b**). While a variety of processes (e.g. translation, gene expression and the nitrogenous compound and peptide metabolic processes) were overrepresented in the deletions that led to growth aberrations in K and Mg, a very specific signature for endosomal transport, vesicle-mediated transport and Golgi-vesicle transport (**Supplementary Figure 4b**, **Supplementary Table 15)** was found for deletion mutants identified to interact with Ca depletion. We also made some unexpected observations. For example, an Adh3 deletion unexpectedly improved the growth rate on Zn depletion media. A potential explanation for this observation is that a deletion of these genes, many of which are likely involved in “zinc-sparing” ^17^ reduces the metabolic cost of synthesis of the proteins they encode (**Figure 4c**).

Next, we also integrated quantitative proteomes of the gene deletions, by filtering a recently published dataset ^24^ to retain deletions of genes bearing metal-related GO annotations. Any protein quantified in these metal-related deletion strains that was differentially abundant *(abs(log_2_(fold difference compared to the control strain)) > log_2_(1.5) and P value (adjusted for multiple testing) < 0.05)* was considered as a significant responder. In total 1391 unique proteins were identified as being differentially expressed when any of the 304 known non-essential metal-related proteins were deleted (**Supplementary Table 16**). The number of proteins identified per metal (1095 for Zn, 548 for Fe, 361 for Mg, 340 for Ca, 187 for Cu, 163 for Mn and 36 for Na) was proportional to the number of annotations for metal binding proteins, with all metals showing an average of six to eight differentially abundant proteins per knockout, except for K and Mn which had 11 and 4 (**Supplementary Table 17**). When we examined the impact of the deletion of genes encoding metal-binding proteins on the abundance of other metal-binding proteins, we found that while most deletions did not significantly alter the abundance of more than 1 or 2 proteins, a few exceptions like *ACO2* and *LEU3* affected the expression levels of numerous metal-binding proteins (**Supplementary Figure 4c**). For calcium and copper-related proteins, a decrease in abundance of metal-related proteins upon the deletion of other metal-related proteins was common. In contrast, iron and zinc-related proteins exhibited mixed responses, indicating complex regulatory interactions within the cell (**Figure 4d**).

Lastly, we included two datasets comprising cellular quantities of metal ions in the *S. cerevisiae* gene deletion and overexpression collections ^25,26,56,59^. For both metallomic datasets, any metal quantity with an absolute *Z-score* as computed by ^27^, *> 1.5* compared to control was considered a significant change. The number of gene deletions that affected a metal ion concentration was quite variable with the highest number identified for Mn, followed by Mo, Na, K, Ca, Zn, Cu and then Fe (**Figure 4b, Supplementary Figure 4d**) while a fairly even distribution of hits across all metals was observed for the overexpression study ^25^ (**Figure 4b, Supplementary Figure 4e**).

These metallomics experiments were conducted using heavy metal supplemented rich YPD media, which is the likely explanation for a high number of hits for Mn and Mo.

When we analysed data from mutants of genes known to bind specific metals, we observed various patterns of cellular metal concentrations. The first was a group of genes encoding Cu, Fe or Mn binding proteins for which a deletion leads to a decrease in cellular quantity of the corresponding metal and an overexpression leads to an increase (e.g., Cu concentration in CUP2 mutants, Mn concentration in YFR006W mutants) (**Figure 4e**). Another group of knockouts displayed an increase in cellular concentration of a metal upon deletion of a gene encoding a protein that binds the same metal (e.g., *CNE1, EDE1, CNB1* (for Ca), *SOD1* (for Cu) and *IDH1* & *IDH2* (for Mg) while displaying a concomitant small increase in the metal or no change in the overexpression mutant. While the metal quantities observed for the first set of genes can be explained by the cell losing capacity to store metal upon deletion and vice versa, the second group likely reflects a disruption of metal homeostasis when regulatory circuits detect a loss of activity of a metal-dependent protein and lead to the upregulation of compensatory mechanisms to take up more of the corresponding metal. For a small minority of genes, exemplified by Cu concentrations in *COX11* mutants, we observed similar cellular metal concentration changes in both knockout and overexpression mutants. Intriguingly, an overexpression of many zinc-binding proteins led to reduced cellular zinc, suggesting the existence of a feedback Zn homeostasis system that modulates zinc uptake, storage, or efflux in response to increased Zn-binding capacity.

### Genetic Interactions and metallomes provide complementary evidence of protein function

The combined dataset provided multiple lines of evidence for the involvement of both, characterised and poorly characterised proteins, in metal ion responses. Moreover, reflecting the orthogonal nature of the datasets, they captured complementary functional properties as well as different sets of proteins. For example, while quantitative proteomics data captures more abundant proteins, which are also likely to be essential, the knout-out strain datasets capture many of the low-abundant proteins. Overall, 1044 ORFs were assessed across all 5 datasets while approximately half the total yeast genome was queried in 3 datasets (**Figure f & g**). In total, 57.6% (3662/6349) of ORFs were associated with a metal ion in at least one dataset (**Figure 4f, left**) with 110 unique ORFs showing a phenotype in at least one metal condition in 3 or more datasets, and only nine showed phenotypes across all 5 datasets (**Figure 4f, left**). When ORF- metal pairs were considered, only two exhibited phenotypes across 4 datasets, 104 across ≥3 datasets, and 1692 across ≥2 datasets (**Figure 4f**, **right**). Notably, both ORFs that exhibited phenotypes across 4 datasets (the nucleolin, YGR159C, which responded to Zn and the Glycogen phosphorylase YPR160W, which responded to K) are predicted to interact with the respective metal ions or common corresponding anions present in metal salts by AlphaFill ^60^.

Thus, proteome, metallome, and genetic interactions provided signals for complementary sets of genes (**Figure 4h**) The proteomics datasets, captured the largest fraction of responses within the known metal-related proteins (**Figure 4i**), followed by the metallomics study of overexpression mutants, metallomics of the deletion mutants and the growth screen of the deletion mutants capturing the lowest signal for known metal-related genes. The highest number of protein associations were found for Zn (676) followed by Fe (316), K (188), Ca (148), Mg (125), Cu (100), Na (81), Mn (45) and Mo (13). All five datasets, combined, assessed 93% (852/914) of all metal-related proteins and 59% (537/914) of these were linked to a metal ion in at least one dataset.

We then turned our attention to poorly characterised genes (**Figures 5a &b**). Understudied proteins for which expression is confirmed (e.g., UniProt annotation score 2), produced a similar number of hits in our datasets compared to well-studied genes (UniProt annotation score >3 (**Figure 5c**)), indicating that our resource could indeed help to mitigate annotation biases. Indeed, 470 poorly characterised proteins, including the aforementioned 55 proteins which were functionally annotated in our ensemble clustering of the proteome, were a hit in at least one of the datasets **Supplementary Table 18 & 19**) We studied two examples in detail. These proteins were identified in different datasets, allowing us to generate hypotheses about their function. The first protein, Ymr196wp, decreases in abundance in conditions with excess Fe (**Figure 5d**). The ensemble clustering pipeline assigned it to Fe cluster 10 based on *metalwise-clustering*, and to cluster 23 in the *allmetal-clustering*. Both clusters were enriched for functional terms related to oxidative stress and chemical stress. Fe cluster 10 was also enriched for proteins that localise to the mitochondria. (**Figure 3a, Supplementary Table 13**). Furthermore, the overexpression of Ymr196wp led to increased cellular Fe concentration (**Figure 5e**), while its protein abundance decreased upon the deletion of seven metal-binding proteins (**Figure 5f**). In parallel, Ymr196wp was also found to associate with the respiratory chain and the TCA cycle, both located to mitochondria, based on proteomic profiles of the *S. cerevisiae* gene deletion collection ^24^. At the metabolic level, the knock-out of the gene results in an accumulation of the amino acid valine ^61^ that requires the mitochondrial enzyme Bat1 ^62^. In summary, Ymr196wp is linked to iron and mitochondrial metabolism based on several independent datasets. Thus, its metal-linked molecular profile suggests that this uncharacterized gene functions in mitochondrial iron homeostasis.

**Figure 5.**
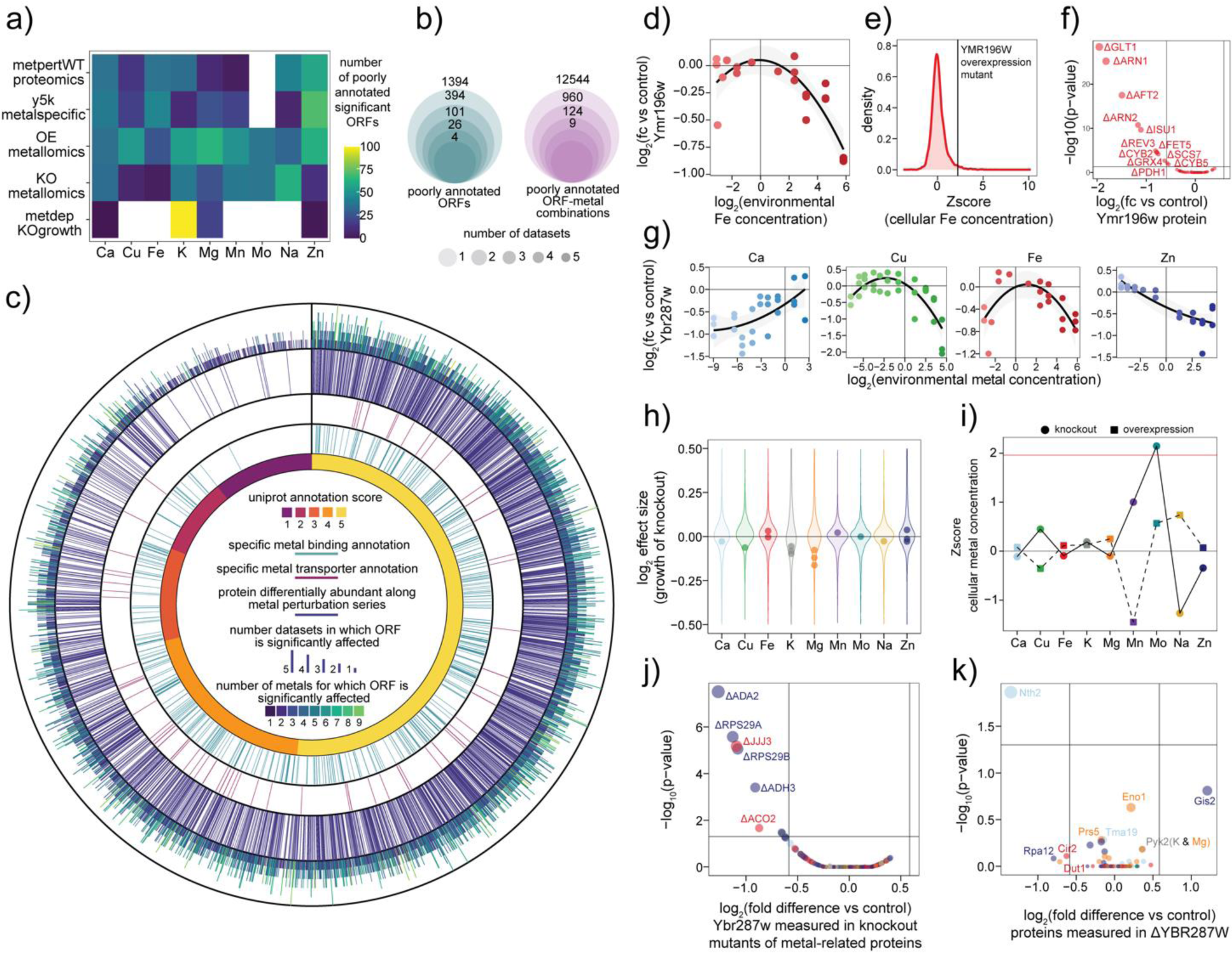
Annotating unknown protein through their metal responsiveness a) *Number of poorly characterised ORFs that were significantly affected in each metal perturbation or metal-related gene deletion (for* ^24^*) across all five datasets.* X-axis: metal that were perturbed or annotated to be connected to the deleted gene. Y-axis: dataset name (refer to Figure 4a for clarity on nomenclature). Colour of each tile indicates the number of poorly characterised ORFs that were significantly affected. Combinations with white tiles represent no significant values. b) *Overlap between significant poorly characterised ORFs (left) and significant ORF-metal combinations (right) across all five datasets.* Size of circle indicates how many datasets are considered with the largest circle representing the set of ORFs or ORF-metal pairs that are significant in any one dataset and the smallest representing those that are significant in all five. Numbers at the top, outside the largest circle represent the total number of unique poorly characterised ORFs and ORF-metal pairs that were measured in all five datasets, cumulatively. c) *Circos plot* ^112^ *depicting the genome-wide scale of all five datasets.* Each index along the circular tracks represents an Open Reading Frame (ORF) in the yeast genome, arranged by UniProt annotation score. Counting from innermost track to outer - the first track indicates the UniProt annotation score (yellow - best characterised proteins with UniProt annotation score of 5 and purple - worst characterised with UniProt annotation score = 1). The second track indicates using ORFs that are annotated as binding to specific metals in the GO-MF database while the third indicates those with specific metal transport annotations. The fourth track indicates ORFs that were significantly differentially abundant in WT *S. cerevisiae* cells cultivated along at least one metal perturbation series in our proteomics dataset. The last, outermost track indicates using colour the number of metals for which an ORF was found to be significantly affect across any of the give datasets and the height of each bar represent the number of datasets in which the ORF was found to be significantly affected (exhibit a gene-metal or protein-metal interaction). d) *Protein abundance of Ymr196w in S. cerevisiae cells upon varying the Fe concentration in cultivation media.* X-axis: log_2_ media concentration of iron relative to synthetic minimal control media, as quantified using ICP-MS. Y-axis: corresponding log_2_(fold difference of Ymr196w protein abundance) e) *Z-score of the cellular Fe concentration of the YMR196W overexpression mutant relative to Z- scores of all other mutants.* X-axis: Z Score cellular Fe concentration. Y-axis: density of mutants at each Z Score. Black vertical line indicates the Z-score of the YMR196W overexpression mutant (2.23). f) *Impact of deletion of iron related proteins on the abundance of Ymr196w protein in each deletion mutant.* X-axis: log_2_(fold difference in protein abundance of Ymr196w in each iron related mutant relative to the *S. cerevisiae* control). Y-axis: -log_10_( P value of significance tests conducted in ^24^ to determine proteins affected in each deletion mutant). Points correspond to Ymr196w protein abundance in each deletion mutant (indicated by gene names on the figure). g) *Protein abundance of Ybr298w in WT S. cerevisiae cells cultivated in each metal concentration series (indicated by panel) along which it was deemed to exhibit a significant change.* X-axis: log_2_(environmental (cultivation media) concentration of each metal relative to synthetic minimal control media) as quantified using ICP-MS. Y-axis: corresponding log_2_(fold difference of Ybr298w protein abundance) h) *Impact of deletion of Ybr298w gene on growth of S. cerevisiae cells in metal depletion media.* X- axis: metal that was depleted. Y-axis: log_2_(effect size (mean colony size of replicates of Ybr287w deletion mutant relative to that of control, divided by the standard deviation of colony sizes of the mutant) - as determined by *pyphe-analyse*) i) *Impact of Ybr298w deletion and overexpression on cellular metal content.* X-axis: metal that was quantified using ICP-MS by ^26^ (knockout deletions) and ^25^ (overexpression). Y-axis: Z-score (calculated in ^27^ of concentration of each metal in the YBR287WW mutants. Colour indicates metal that was quantified. Shape indicates the type of mutation: circles indicate knockout deletion and squares indicate overexpression. j) *Impact of deletion of metal related proteins on the abundance of Ybr298w protein in each deletion mutant.* X-axis: log_2_(fold difference in protein abundance of Ybr298w in each metal related mutant relative to the WT *S. cerevisiae* control). Y-axis: -log_10_( P value of significance tests conducted in ^24^ to determine proteins affected in each deletion mutant). Points correspond to Ybr298w protein abundance in each deletion mutant (indicated by gene names on the figure). Colours indicate the metal annotation of the deleted gene based on the GO-MF database. k) *Impact of the deletion of Ybr298w gene on the abundance of metal-related proteins.* X-axis: log_2_(fold difference in protein abundance of different proteins in the Ybr298w mutant relative to the WT *S. cerevisiae* control). Y-axis: -log_10_( P value of significance tests conducted in ^24^ to determine proteins affected in the Ybr298w deletion mutant). Points correspond to protein abundance of each metal-related protein (indicated by gene names next to points) in the Ybr298w deletion mutant Colours indicate the metal annotation of the measured protein.

Our second example, Ybr287wp localises to the endoplasmic reticulum ^63^ and contains eight transmembrane domains ^64,65^. The abundance of Ybr287wp was correlated to environmental Ca, Cu, Fe and Zn concentrations (**Figure 5g**) and it was associated to a cluster enriched in iron transport and cation channel terms (cluster 14, see **Figure 3b**). Deletion of YBR287W decreased growth rates in K and Mg depletion and led to milder changes in Cu, Fe, Mn and Na depletions (**Figure 5h**). The knockout and overexpression mutants revealed mirrored cellular metal concentration profiles with altered cellular concentrations of Cu, Mn, Mo and Na and minor alterations in Ca, Fe and Zn (**Figure 5i**). Furthermore, the deletion of eight other metal-binding proteins led to downregulation of Ybr287wp (**Figure 5j**) while the deletion of Ybr287w itself led to a significant decrease in the abundance of the Ca-binding protein Nth2 and smaller changes in Zn and Mg binding proteins (**Figure 5k**). Collectively, Ybr287wp is associated with metal biology based on several datasets. The molecular profile of this uncharacterized transmembrane protein is consistent with that of a promiscuous metal ion transporter.

### Metal dependency of a subset of highly connected metabolic enzymes translates to network-wide metal responsiveness

Metabolism was identified by our functional enrichment analyses as one of the cellular networks most affected by metal availability. Therefore, we selected the metabolic network of *S. cerevisiae* to exemplify the utility of our dataset to assess system-wide impact and implications of metal availability. Gene Ontology annotations suggest that 26% of enzymes and ∼13% reactions in the genome scale metabolic model Yeast8 (**Supplementary Table 20**), 29% of enzymes in the Enzyme Commission (EC) database and 17% of those in the KEGG database, are linked to at least one metal through direct binding, transport, or complex metal-containing-cofactor binding (**Figure 6a, left** and **Figure 6b)**. Furthermore, 76% of all Enzyme Commission (EC) numbers and 89% of all KEGG pathways involve at least one metal-associated protein (**Figure 6a, right**).

**Figure 6.**
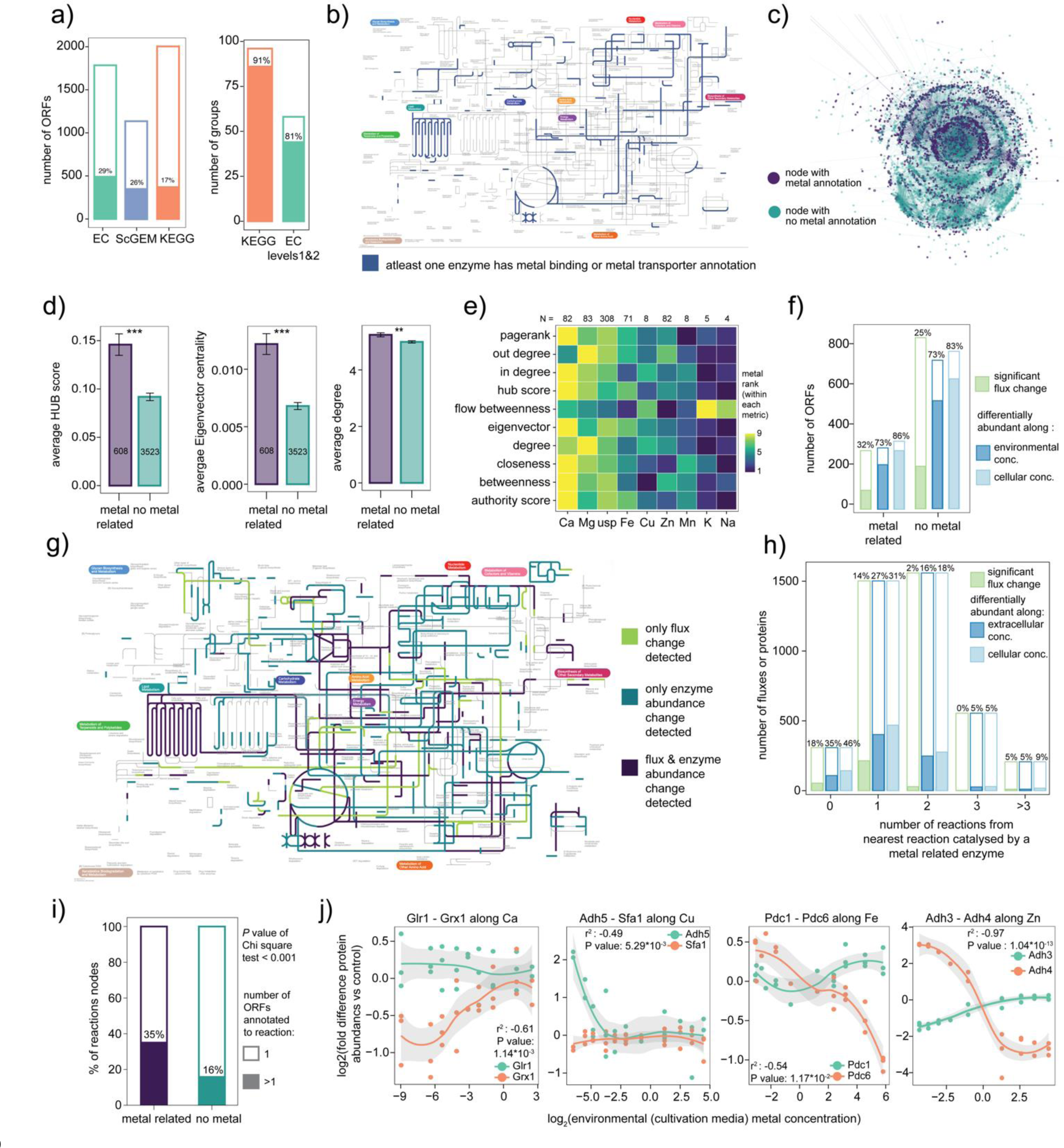
Central role of metal-dependent enzymatic reactions results in a network-wide metal dependency of metabolism a) *Fraction of enzymes in the enzyme commission & KEGG databases and the Yeast8 genome scale metabolic model* ^113^*) that are annotated to be connected to metals.* Left panel summarises annotations at the ORF level. X-axis: database (EC: Enzyme Commission numbers ^114^, ScGEM : Yeast8 genome scale metabolic model ^113^ and KEGG : Kyoto Encyclopedia of Genes and Genomes ^115^). Y-axis: outer rectangle of bar indicates total number of ORFs in the database and inner (filler) bar indicates the number of ORFs that have any metal-related annotations in the GO-MF database. Numbers indicate the percentage of ORFs with metal related functions in each database. Colours are used to differentiate databases. Right panel summarises annotations at the pathway and enzyme class (two levels of the enzyme commission numbers) level. X-axis for right panel: database. Y-axis: outer unfilled rectangle of the bar plot indicates the number of pathways (for KEGG) and the number of unique EC numbers up to level 2 that were considered while the inner filled bar represents the number of KEGG pathways or EC number categories for which at least one member ORF had a metal-related annotation. Numbers in each bar indicate the percentage of KEGG or EC categories that contained at least one ORF with a metal-related annotation. b) *Metal-related enzymes in the S. cerevisiae metabolic network.* Map of the *S. cerevisiae* metabolic network with blue colour indicating reactions that are catalysed by at least one metal- related enzyme (there can be multiple enzymes assigned to one reaction). Created using iPATHv3 ^110^. c) *Visualisation of the directed bi-partite graph representation of the S. cerevisiae Yeast8 GEM*. Each metabolite and each reaction is represented by a unique node in the graph. Links in the graph are directed (from substrate metabolite to reaction and from reaction to product) with irreversible reactions added twice (with links in both directions). The graph was created using the python *igraph* library and visualised using the Kamada Kawai method. Colours indicate whether each node has a metal-related annotation or not. Reaction nodes that map to at least one ORF with a metal-related annotation are marked purple and metabolite nodes with even one direct link to a reaction node that is annotated as metal-related are also marked purple. All other metabolite and reaction nodes are marked teal. d) *Metal related reaction nodes in the S. cerevisiae metabolic network are more central and connected compared to nodes without metal annotations*. Each panel represents a different metric (HUB score, Eigenvector centrality and degree). X-axis: metal annotation status. Y-axis: bars indicating the mean (bar height) of each metric in each category. Error bars represent ± standard error around the mean. Three stars above the bars represents a P value of < 0.001 and two stars represent P value < 0.01 based on t-tests conducted between the two groups (metal related annotation and no metal related annotation) for each metric. e) *Centrality metrics of reaction nodes annotated to bind each metal.* X-axis: metal annotation of a reaction node (‘usp’: unspecific annotation which indicates that an ORF mapping to the reaction node is annotated as having a metal binding or other connection to metals, but it is unclear which metal it is connected to). Y-axis: metric that was calculated using the python *igraph* library. Colour indicates the rank of each metal within each metric. Numbers at the top of the tile plot indicate the total number of reaction nodes that were used for the calculation for each metal. f) *Fraction of the metabolic network that responds at the flux (simulated) and protein abundance (experimentally quantified) level to perturbations of metal availability.* X-axis: metal annotation status. Y-axis: outer unfilled rectangle of bars represents the total number of enzymes measured or fluxes assessed in the simulation, inner filled bar represents the number of enzymes for which a significant flux change through the enzyme was detected in the simulation (light green), protein abundance change was detected in linear models fit along environmental (cultivation media) metal concentration (dark blue) or cellular metal concentration (light blue) for at least one metal. Numbers above each bar indicate the percentage of fluxes or protein abundances that were significantly altered. g) *Metal responsiveness of the S. cerevisiae metabolic network.* Light green colour indicating reactions through which a significant flux change was detected. Teal indicates those for which only enzyme abundance changes were detected at the protein level (based on linear models along either environmental (cultivation media) concentrations or measured cellular concentrations. Purple indicates reactions for which both the flux simulations and the experimental protein abundance data indicate a significant change. Created using iPATHv3 ^110^. h) *The shortest distance to a metal-related node correlates with the likelihood that any node in the metabolic network will be affected by perturbed metal availability.* X-axis: shortest distance in number of reactions from the nearest metal-related node (as calculated using *igraph*). Zero indicates that the reaction node itself maps to at least one ORF with a metal-related annotation. Y-axis: outer unfilled rectangle of bars represents the total number of enzymes measured or fluxes assessed in the simulation, inner filled bar represents the number of enzymes for which a significant flux change through the enzyme was detected in the simulation (light green), protein abundance change was detected in linear models fit along environmental (cultivation media) metal concentration (dark blue) or cellular metal concentration (light blue) for at least one metal. Numbers above each bar indicate the percentage of fluxes or protein abundances that were significantly altered at each distance from the nearest metal-related reaction node. i) *Metal-related reaction nodes are more likely to bear multiple enzyme annotations compared to those without metal-related annotations.* X-axis: metal annotation status. Y-axis: outer unfilled rectangle of bars represents the total number of reaction nodes that were assessed in each group, inner filled bar represents the number of reaction nodes which had more than one enzyme annotation in the Yeast8 metabolic model. Numbers represent the percentage of nodes in each category that have more than one enzyme annotation. The difference between the two groups based on a Chi-square test was significant with a P value < 0.001. j) *Selected examples of enzyme-pairs mapping to metal-related reaction nodes that exhibit negative correlations in protein abundance*. Each panel represents a different enzyme-pair indicated at the top of the panel. X-axis: log_2_(environmental (cultivation media) metal concentration). Y-axis: log_2_(fold difference in protein abundance in cells cultivated in metal perturbation condition relative to those cultivated in synthetic minimal control media). Colour is used to differentiate between the two enzymes in each pair and have no shared meaning across panels.

Oxidative phosphorylation bears the strongest relationship to metalloenzymes (80%), closely followed by folate biosynthesis (75%) (**Supplementary Table 21**). To conduct network analysis, we represented the Yeast8 genome-scale metabolic reconstruction as a directed, bipartite graph (**Figure 6c**). We revealed that nodes with metal-related annotations occupied more central positions, as evidenced by multiple centrality metrics such as HUB score, Eigenvector centrality, and degree (**Figure 6d & Figure 6e**). As a consequence, even though only 13% of the reactions directly involve a metal ion, 43% of the reaction nodes are only one reaction away from a metal dependent enzyme (**Supplementary Table 20**). At the metabolite level, a staggering ∼71% of metabolites were either directly connected or one reaction away from an enzyme with metal-related annotation (**Supplementary Table 22**).

We next simulated flux level responses using the CofactorYeast framework ^66^ (**Methods)** and compared these to the experimentally observed enzyme level responses. Simulated flux-level responses were lower (32% of simulated reaction fluxes for metal-related enzymes and 25% for non-metal-annotated enzymes were changed by more than 50%) than those experimentally observed (∼73% of both metal-related and non-metal-annotated enzymes) (**Figure 6f, 6g)**. At the per-metal level, Zn perturbations elicited the highest response at both levels with ∼19% of fluxes and ∼49% of enzymes being affected (**Supplementary Figure 5a**, **Supplementary Table 23**).

Interestingly, enzymes without pre-existing metal annotations in the GO database had similar response levels as those without (**Figure 6f, Supplementary Table 23**). Enzyme responsiveness was closely linked to the proximity of the reaction node to reactions catalysed by one or more metal-requiring enzymes. Reactions catalysed by metal-binding enzymes were the most affected by metal perturbation, with 18% of fluxes and 35% of enzyme levels showing changes in abundance due to environmental perturbations, and 46% responding to variations in cellular metal concentrations. Enzymes more than two reactions away from metal-requiring reaction nodes showed minimal impact (**Figure 6h).**

The relatively low flux-level impact compared to the enzyme-level impact suggested that metabolic fluxes could potentially be buffered against loss of essential metal availability. One potential mechanism is the presence of isoenzymes with differential metal requirements. To test this possibility, we extracted all reactions with more than one associated ORF from the genome scale metabolic model (**Methods**), which include multi enzyme complexes. We found that metal related reactions were more likely to be catalysed by more than one isozyme (**Figure 6i)**.

Furthermore, of the 246 unique combinations of metal-binding enzyme pairs in which at least one enzyme was differentially expressed, 43 instances corresponded to enzyme pairs that respond in an anti-correlated manner along a metal concentration series, i.e., we observed a simultaneous increase in the abundance of one isoenzyme and a decrease in abundance of the other or vice versa (**Supplementary Table 24**). An illustrative example is that of Adh3 and Adh4, both of which are annotated to bind Zn^2+^ with Adh4 suggested to potentially bind Fe^2+^ under Zn limitation ^67^. The protein abundance patterns corroborate the hypothesis, the Fe^2+^ dependent Adh4, is inducted upon Zn limitation conditions (**Figure 6j** Pearson correlation coefficient = -0.97 and *P* value = 1.04*10^-13^) and decreases in abundance in Fe limitation (**Supplementary Figure 5b**). Twenty eight such isozyme pairs were identified along the Zn perturbation series, another 15 each along Ca, Cu, Fe and Na perturbation gradients with 11 pairs being negatively correlated along more than one metal perturbation series (**Supplementary Table 24**).

## Discussion

The critical role of metal ions in cellular biochemistry dates to the origin and evolution of metabolism. In the low-oxygen atmosphere of early Earth, iron existed in its reduced, water-soluble form as Fe(II). The high concentration of Fe(II) found in Archean sediments suggests it was present in significant amounts in the world’s early oceans ^68,69^. Consequently, Fe(II) was readily available both as an electron donor and as a catalyst for the evolving metabolic network. Recent experiments have demonstrated that Fe(II) can catalyse non-enzymatic interconversion that closely resembles central metabolic pathways, such as glycolysis, pentose phosphate pathway, TCA cycle and cofactor biosynthesis, suggesting that the structure of these central metabolic pathways was shaped by metal-catalysed chemistry ^70–74, 75, 76^. In addition to the reliance of the core metabolic network on metal-catalysed reactions, the development of new protein domains throughout the evolutionary history of life was likely influenced by changes in the availability of metals on Earth’s surface. ^2,77^.

The availability of metal ions can vary across a range of time and length scales such as evolutionary periods, geological landscapes and ecological niches. Combined with the importance of metal ions for metabolic and protein function, this variability in metal availability has therefore led to the evolution of sensing, transport, and buffering systems for metal ions.

Specifically in well-studied eukaryotic cells, many transporters, chaperones, and metal-responsive transcriptional elements that maintain metal ion balance have been discovered ^7,10^. However, our understanding of how biological networks respond to the physiologically relevant changes in metal availability on a broader scale, signalling systems contributing to metal ion homeostasis and their connection to cellular phenotypes remains surprisingly limited.

In this study, we aim to address the substantial gap in understanding of cellular responses to fluctuations in metal ion availability, by addressing two key contributing factors. First, we noticed that metal ion availability has thus far been modified within the physiological (non-toxic, non-limiting) concentration range only in a minuscule fraction of molecular biology and systems biology experiments. Consequently, the responsiveness of cellular networks to changes in metal ions is understudied. Second, cellular metal concentrations do not necessarily mirror environmental (media) levels due to the capacity of cells to buffer against environmental fluctuations and the promiscuity of metal ion transport systems which results in interactions between metal ion concentrations. To address the first limitation, we varied all major metallic components of cultivation media over five orders of magnitude in concentration. Since synthetic growth media for *S. cerevisiae* have been well-defined since the 1950s ^28,78^, it allowed us to vary each metal ion concentration typically supplied in minimal media. To address the second, we apply quantitative metallomics, to contrast cellular and environmental metal ion concentrations, as well as to systematically detect interactions between the concentration series. To achieve this, we adopted an extensively validated protocol ^26^ for metallomics sample preparation combined with inductively coupled plasma mass spectrometry (ICP-MS) to distinguish between environmental (media) and cellular concentrations of the metal ions.

We selected *S. cerevisiae* for this investigation because it is an extensively studied model organism, especially in the context of metal ion biology. This allowed us to leverage prior knowledge to assess responses of the large number of proteins bearing metal-binding, metal- transporters, and other curated gene function annotations, and integrate genome-scale datasets such as genome-scale metallomic profiles ^25,26^, proteomes of a genome-scale knock-out collection ^24^ and a genome-scale metabolic model that includes metal ions as cofactors ^66^ that are thus far uniquely available for yeast. In addition to its scale, a key factor that distinguishes our study from previous work addressing metal biology systematically, is that we utilised prototrophic strains and minimal media formulations that lack amino acid supplements. As a result, our experimental setup takes into account that one of the primary functions of metal ions is as cofactors in enzymes that catalyse key reactions in biosynthetic metabolism, many of which are feedback inhibited in rich growth media ^79^. These combined efforts yielded a much more comprehensive picture about the role of the metal ion concentrations in cellular networks and revealed a remarkable interdependence of cellular processes and their metallomic environment, at the scale of the proteome.

While our dataset is intended to serve as a resource for the research community to study metal ion biology at various molecular layers, we derive several general principles that govern cellular responsiveness to metal ion perturbations. For example, in addition to providing a more systematic and quantitative perspective on concentration-buffering of metal ions against extracellular fluctuations, we reveal that metal ion homeostasis strongly varies between metals and is evident only for those that are physiologically important. For instance, molybdenum was not buffered and elicited cellular responses only at high, toxic concentrations, while its depletion caused no growth defects or protein responses. We thus conclude that *S. cerevisiae* cells do not require molybdenum. This suggests that the decades-old practice of supplementing molybdenum to all common yeast media formulations should be reconsidered. Our results also indicate that concentrations of most essential metals in synthetic minimal media are well in excess of those required for normal cell growth, supporting earlier conclusions ^78^. While this practice is not inherently problematic - we find that routinely supplied concentrations of essential metals well within the physiological, non-toxic range - it does imply that relevant phenotypes, such as ion transport defects, might be masked in experiments conducted in these media. Therefore, lowering the concentration of metal ions compared to the standard media recipes might lead to new discoveries. In parallel, we report comprehensive quantitative data about the interdependence of cellular concentrations of several metals. These results help not only the interpretation of our own data, such as the proteomes, but could also provide key context for the interpretation of results of other metal ion perturbation experiments.

Our study provides a comprehensive blueprint for understanding how cells adapt to variations in metal ion concentrations at the molecular level. The systematic nature of the data unveils a comprehensive cellular response to changes in metal availability and highlights how these responses are integrated across different layers of the cell. Even though our experiments addressed the response to metal concentration changes in a single environmental condition and within a single genetic background of one eukaryotic species, we discovered that the abundance of approximately 60% of proteins is influenced by metal availability, with zinc, iron, calcium, and copper eliciting the most profound responses. We can thus speculate that many biological responses reported in the literature, will be dependent on the metal ion levels available to cells. In this context it is interesting that major components of the cellular transcription and signalling machinery, especially most kinase pathways, are among the metal-responsive pathways. For example, we reveal that proteins in 28 out of 34 signalling pathways that are captured by our proteomes, change in abundance to several metal ions. These pathways include mTOR^78^, a signalling pathway that functions at the crossroads of cellular transport, the lysosome, and energy metabolism, all processes related to metal biology. Notably, we can exclude changes in growth rate as a main driver for most of these responses: proteomic responses explained by a change in growth rate were detected in the case of zinc and potassium depletion, and under conditions with extremely high levels of iron and copper. Thus, our study reveals that metal ion responsiveness is an underappreciated aspect of cellular regulation and signalling.

Third, our resource provides a fresh perspective about a common problem in molecular biology - the high number of understudied proteins Even in the most well-studied organisms, many proteins lack functional annotation. We speculated previously that a limited number of experimental conditions tested in laboratory experiments might be a leading cause of missing protein function ^81^. Our study supports this argument. We find that understudied proteins, except for those for which expression has not been confirmed, are as likely to be a hit across our datasets, as proteins with high annotation scores. Systematically varying metal ion levels can thus help to mitigate the annotation bias and provide testable hypotheses about new protein function. We highlight two examples that are linked to metal biology across multiple datasets These are Ymr196w, which bears a molecular profile consistent with being involved in iron homeostasis, and Ybr287w, which has a profile of a promiscuous metal ion transporter.

Lastly, we use cellular metabolism as an example of a network that is impacted by metal availability. We find that even though metal-dependent reaction nodes comprise a moderate ∼13% of the metabolic network, the central location of these reactions leads to a high metal responsiveness at the enzyme abundance level and a staggering ∼70% of metabolites are only one reaction away from metal-dependent reactions. We speculate that the centrality of metal- related nodes stems from the key role of metal ion catalysis in early metabolic evolution and that the evolution of metal requiring enzymes could be more constrained relative to other proteins due to the essential catalysis they enable. Moreover, we report that metal-dependent reactions are more likely to be catalysed by isozymes, and a subset of these involve enzymes that catalyse similar reactions but use different metal cofactors. We speculate that the central role of metalloenzymes, combined with dramatic changes in metal availability across time and ecological niche, was one of the drivers of divergent enzyme evolution.

## Implications of our study

Our comprehensive resource illuminates the pivotal role of metal ion homeostasis within the regulatory and functional landscape of the cell, promising to redefine our understanding of cellular processes. We envision this dataset serving as a foundational reference for unravelling the connections between metal ions and the spectrum of biological processes, facilitating the integration of metal ions into a system wide understanding of cellular function. This tool opens avenues for exploring the roles of previously understudied genes, enriching our comprehension of signalling pathways, and gene regulatory networks.

Furthermore, our findings advocate for a paradigm shift in current laboratory practices, which starkly contrast the dynamic metal ion concentrations found in natural settings. Typically, experimental conditions do not account for the natural variation in metal ions and fall short of reporting of metal ion levels. While the former may mask biological discoveries, the latter potentially hampers the reproducibility of laboratory research. By varying and diligently reporting metal ion concentrations, researchers can unlock new biological insights and enhance experiment reproducibility.

## Limitations

Despite the systems-scale and quantitative nature of all our experiments, our study has limitations that need to be considered while interpreting and querying our dataset. The first is that we restricted the scope of our study to a single species and a single reference growth condition. The biological response to metal ion perturbation is therefore likely to be more extensive than that reported herein. Second, we measure total cellular metal concentrations and do not address differences in metal ion concentrations between subcellular compartments which can vary significantly and are known to impact protein function within compartments. Future studies could build on our work and assess the impact of subcellular metal distribution and resolve cellular responses to perturbed metal availability at the subcellular scale. A further limitation are the analytical constraints we faced while varying and quantifying certain metals. The concentration range across which potassium and magnesium could be varied was limited and our ICP-MS could not accurately quantify copper and molybdenum at levels exceeding those in standard synthetic minimal media. Therefore, it is likely that we have missed out on the metal responsiveness of some genes, pathways, and processes. Lastly, while our findings lend strong support to many proposed functions of poorly characterised proteins, the mutant libraries used to validate our hypotheses suffer from limitations common to such libraries, such as secondary mutations. Hence, we advocate for the use of our resource to derive system-level insights into the role of metals in biology and as a foundation for hypothesis generation to be validated by future studies.

## Methods

### EXPERIMENTAL MODEL DETAILS

#### Strains and mutant libraries

*Saccharomyces cerevisiae* (S288C) haploid (*MAT*a) was used as the experimental model system. Specifically, the S288c derivate BY4741-kanmx4::*his3* rendered prototrophic with the pHLUM minichromosome ^29^ was used as the wild type (WT) strain for the growth rate, cellular metallomics and proteomics experiments in metal perturbation media conditions. The strain was chosen for consistency with previous work, in which we determined proteomes and amino acid metabolomes for the knock-outs ^1,2^, allowing us to directly compared the datasets. The same knockout mutant library ^29^ was employed for the growth screen under metal depletion conditions on agarose media.

#### Cultivation of WT cells

The WT *S. cerevisiae* cells were revived from cryostocks on YPD (Yeast Peptone Dextrose) agar plates and incubated at 30°C for ∼24hrs (until colonies appeared). A single colony was picked and streaked onto SM (Synthetic Minimal media) agarose plates and incubated at 30°C for ∼36 hrs (until colonies appeared). Then, a single colony from the SM plate was used to inoculate the 5mL starter liquid SM culture which was incubated on a shaker at 30°C for ∼36 hrs. The pellet from this culture was washed three times with water and resuspended in SM0 media (synthetic minimal media without addition of metal salts except for KH_2_PO_4_, Mg_2_SO_4_ and ZnSO_4_.7H_2_O, detailed composition in **Supplementary Table M1**.). The resuspended pellet was used to inoculate 300mL of SM0 media such that the OD_600_ of the culture at inoculation was 0.05. After incubation in a shaker at 30°C, the pellet from SM0 culture was washed three times with water and used to inoculate deep-well 96-well plates (Eppendorf, 10052143) filled with each of the 91 cultivation media containing perturbed metal concentrations (see **Supplementary Table M1** for detailed media composition) such that the starting OD_600_ of the culture was 0.05. Four replicates of each cultivation condition were distributed across at least two different 96-well plate layouts (**Supplementary Table M2**). Each 96-well plate contained six control media (identical to SM but created in the same manner as all other media and termed Allele media in supplementary tables). The borders of each plate were filled with water instead of cultures to avoid edge effects that were observed in preliminary tests. Three wells in each plate were emptied and filled with technical controls for mass-spectrometry measurements post-cultivation. All media metal perturbation media were prepared in plastic, all reagents used for preparation were of ICP-MS grade, except for glucose, and only deionized water that had previously been checked for metal contamination on the ICP-MS was used for preparing media or washing the cells. All deep-well 96-well plates were covered with Breathe-Easy seals during cultivation.

#### ICP-MS measurements of cultivation media

All cultivation media bearing variations in metal concentrations were analysed using ICP-MS to quantify the concentration of each metal. A 17-point calibration series was prepared fresh (up to 24 hours before measurement) using the certified metal standards (see **Key Resources Table** ). Details of the concentration of each element in each calibration standard are available in **Supplementary Table M3** and plate layouts for ICP-MS measurements of media are in **Supplementary Table M4**. All samples were measured on an Agilent 7900 ICP-MS coupled to an SPS-4 auto-sampler and an Agilent MicroMist nebulizer. The instrument was operated with Nickel (Ni) cones and the measurement parameters were optimised using the Tuning solution and the PA solution. The following gas modes were used for different metals: Helium (^24^Mg, ^59^Co, ^63^Cu, ^66^Zn), High Energy Helium (^32^S, ^31^P, ^55^Mn), and Hydrogen (^39^K, ^40^Ca, ^56^Fe) mode. Details of all peristaltic pump settings, tune parameters and all raw data from the ICP-MS can be found in **Supplementary Note M1**). Twenty individual media had incorrect concentration so the perturbed metal and were therefore remade and re-measured before processing with inoculation of yeast cells for growth, metallomics and proteomics characterisation.

#### Growth rate measurement of WT cells

WT *S. cerevisiae* cells were prepared as described above and inoculated into short-well 96-well plates with 180μL of media in each well. These plates were then placed in a Spark-Stacker (TECAN) plate reader operating in kinetic mode and sequential acquisition of multiple 96-well plates. Absorbance at OD_600_ was calculated from the mean values of five multi-well reads obtained every 30 minutes for 48 hours for each position.

#### ICP-MS measurements of *S. cerevisiae* cells

Media to be used for cultivating cells for intracellular metal quantification with ICP-MS was through a PVDF membrane plate (Agilent 200931-100) into fresh deep-well (2mL) 96-well plates immediately before cells were inoculated into it to ensure no insoluble precipitates remain in the media that could interfere with washing of cells on PVDF membranes post-cultivation and remain in the cell digest to be used for ICP-MS measurements. WT *S. cerevisiae* cells were prepared, inoculated into deep-well 96-well plates (Eppendorf, 10052143) containing metal perturbation media and cultivated on a shaker at 30°C for 24 hours as described above. After cultivation, cells were collected by filtering cultures in each deep-well 96-well plate through a 96-well PVDF membrane plate. Yeast cells on the 96-well PVDF membranes were then washed 3-times with a solution composed of 10μM EDTA and 3μM TrisHCL. Centrifugation speeds and durations had to be modified on the go to ensure the entire volume of cells in each culture passed through the PVDF membrane. PVDF membrane plates bearing the washed cells were incubated on a hot plate at 70°C until completely dry. Internal metal standards were added to the membranes and the dried cell pellet was then digested by adding 60μL of HNO_3_ and heating at 94°C for 40 minutes. Deionised water was added to each digested pellet to achieve a final HNO_3_ concentration of 10% (v/v). Due to small differences in evaporation rates, slightly different volumes remained in each plate after incubation at 94°C. Therefore, the total amount of deionized water to be added and the final volume available for ICP-MS varied and is noted in **Supplementary Table M5.** The diluted cell extracts from one batch (96-well plate) at a time along with the fresh calibrants were measured on an Agilent 7900 ICP-MS coupled to an SPS-4 auto-sampler and an Agilent MicroMist nebulizer using the same methodology as described above in ‘**ICP-MS measurements of cultivation media’.**

#### Proteomics sample preparation

*S. cerevisiae* cells were prepared, inoculated into deep-well 96-well plates (Eppendorf, 10052143) (Eppendorf, 10052143) containing metal perturbation media and cultivated on 30°C with 1000 rpm shaking (Heidolph Titramax incubator) for 24 hours as described above. The methodology used for peptide extract preparation from cell pellets and measurement of mass spectrometry data was identical to that reported in ^24^. Briefly, after cultivation, 50uL of culture was removed from each well and transferred into a transparent short-well 96-well plate pre-filled with 50uL H_2_O and OD_600_ measurements were recorded. Each deep-well plate was centrifuged at 3220 rcf (Eppendorf Centrifuge 5810R) to pellet the cells, Breathe-Easy seals and the supernatant were removed, and the plates were sealed with aluminium foil (adhesive PCR plate foil) and a plastic lid and frozen at -80°C until further processing. Protein extraction and digestion was carried out in batches of 4 96-well plates (384 samples). To reduce batch effects, stock solutions (120 mM iodoacetamide, 55 mM DL-dithiothreitol, 9 μl 0.1 mg/ml trypsin, 2 μl 4x iRT) were prepared in one batch and stored at –80°C until required. Stock solutions of 7 M urea, 0.1 M ammonium bicarbonate, 10% formic acid were stored at 4°C. All pipetting steps were carried out with a Beckman Coulter Biomek NX^P^ liquid-handling robot.

To lyse the cells, 200 μl 7 M urea / 100 mM ammonium bicarbonate and glass beads (∼100 mg/well, 425–600 μm) were added to the frozen pellet. Then, each plate was sealed with a silicone mat (Cap mats, (Spex) 2201) and cells were lysed using a Geno-Grinder (Spex) bead beater for 5 min at 1,500 rpm. After centrifuging the plates for 1 minute at 4,000 rpm, 20 μl 55 mM dithiothreitol (DTT) was added and mixed to achieve a final concentration 5 mM DL). The samples were then incubated for 1 h at 30°C after which 20 μl 120 mM iodoacetamide was added (final concentration 10 mM) to each well. The plates were incubated for 30 min in the dark at room temperature before adding 1 mL of 100 mM ammonium bicarbonate. Each plate was centrifuged for 3 min at 4,000 rpm and then 230 μL of the supernatant was transferred to prefilled trypsin plates. The samples were incubated at 17h at 37°C for trypsin digestion after which 24 μl 10% formic acid was added to each well. The digestion mixtures were cleaned up using BioPureSPN PROTO C18 MACRO 96-well plates. For solid-phase extraction, samples were centrifuged for 1 min at various speeds (listed below) using an Eppendorf 5810R centrifuge 5810R. For the solid-phase extraction, each plate was conditioned with methanol (200 μl, centrifuged at 50 *g*), washed twice with 50% ACN (200 μl, centrifuged at 50 *g* and flow-through discarded), equilibrated three times with 3% ACN, 0.1% FA (200 μl, centrifuged at 50 *g*, 80 *g*, 100 *g*, respectively, flow-through discarded). Finally, 200 μl of each trypsin digested sample was loaded onto the solid phase extraction column plates, centrifuged at 100 *g* and washed three times with 200uL of 3% ACN & 0.1% FA solution (centrifuged at 100 *g*). After the last washing step, the plates were centrifuged once more at 180 *g* before the peptides were eluted in 3 steps (eluted twice with 120 μL and one with 130 μL 50% ACN, centrifugation at 180 *g*) into a collection plate (1.1 mL, V-bottom). Collected material was completely dried in a vacuum concentrator (Concentrator Plus (Eppendorf)) and redissolved in 40 μL of 3% ACN & 0.1% formic acid solution before being transferred into a 96-well plate (700 μL round, Waters, 186005837) prefilled with iRT peptides (2 μL, diluted to 1:32). Quality control samples for repeat injections were prepared by pooling digested and cleaned-up control samples from all the 96-well plates. To quantify total peptide concentration, 2 μl of each sample were loaded onto Lunatic microfluidic 96-well plates (Unchained Labs). Peptide concentrations were measured with the Lunatic instrument (Unchained Labs). Total peptide concentration in each peptide extract was calculated from the absorbance value at 280 nm and the protein-specific extinction coefficient.

#### Liquid chromatography–mass spectrometry

The digested peptides were analysed on a nanoAcquity (Waters) running as microflow LC (5 μl/min), coupled to a TripleTOF 6600 (SCIEX). 2 μg of the yeast digest (injection volume was adjusted for each sample based on the measured peptide concentration) were injected and the peptides were separated in a 19-min nonlinear gradient (**Supplementary Table M8**) ramping from 3% B to 40% B (solvent A: 1% acetonitrile/0.1% formic acid; solvent B: acetonitrile/0.1% formic acid). A HSS T3 column (Waters, 150 mm × 300 μm, 1.8 μm particles) was used with a column temperature of 35°C. The DIA acquisition method consisted of an MS1 scan from m/z 400 to 1250 (50 ms accumulation time) and 40 MS2 scans (35 ms accumulation time) with variable precursor isolation width covering the mass range from m/z 400 to 1250 (Table S2).

Rolling collision energy (default slope and intercept) with a collision energy spread of 15 V was used. A DuoSpray ion source was used with ion source gas 1 (nebuliser gas), ion source gas 2 (heater gas), and curtain gas set to 15 psi, 20 psi, and 25 psi. The source temperature was set to 0°C and the ion-spray voltage to 5,500 V. The measurements were conducted over a period of 2 months on the same instrument.

#### Growth screen of knock-out deletion mutants

To explore the contribution of each non-essential gene to fitness on media depleted of each metal, we performed a growth assay. The prototrophic *S. cerevisiae* haploid knock-out collection (PHKo) ^29^ was grown on 20 different media (corresponding to depletion of the metals Ca, Cu, Fe, K, Mg, Mn, Mo, Na and Zn and the control, for detailed list and composition see **Supplementary Table M9**) chosen after pre-tests in which we cultivated WT prototrophic *S. cerevisiae* BY4741+pHLUM strain on agarose media containing various concentrations of metal salts. These include two types of depletion for Ca and Fe (Ca omission, Ca omission with chelator EGTA (Ethylene glycol tetraacetic acid), Fe omission and Fe omission combined with the chelator dipyridyl (DiP)), three concentrations of K, Mg and Zn and three types of controls (synthetic minimal (SM) media, SM with DiP and SM with EGTA). The PHKo library was revived from YPD (Yeast extract Peptone Dextrose) + glycerol stocks in 96-well plates frozen at -80°C on YPD-agar and then combined with a grid of the control strain for the library (BY4741+pHLUM *his3Δ*) into a 1536 spot layout on SM-agarose in 4 different arrangements. Thus, our assay contains 4 biological replicates of each strain in the PHKo collection. Since the growth of neighbouring strains may affect the colony size of a strain, our re-arrangement strategy allows us to consider neighbourhood effects on colony size before making inter-strain comparisons. For the reference grid, control strain BY4741+pHLUM *his3Δ* was streaked out on SM agar and grown at 30°C for 2 days. A 24h culture of a single colony was made in 40 mL of liquid YPD media and pinned from a bath on YPD agar in 96-spot format and incubated at 30°C for 2 days. The PHKo library was revived from cryostocks in 384 format on YPD agar and incubated for two days at 30°C. A custom Singer ROTOR HD^TM^ program was used to reshuffle the library (using 96 short pins) into 4 random arrangements, consisting of 5 plates each, on standard SM media. At this stage, the reference grid was combined with the library by pinning the reference strain colonies onto the combined plates into the A1, D4 and C2 sub-positions in 1536 format. These combined plates, containing all the library strains and the reference grid were cultivated at 30°C for 2 days and then copied onto fresh SM agarose plates to obtain a clean source plate with evenly spaced and sized colonies. Three copies of each combined plate were made, yielding 60 source plates in total which were incubated at 30°C for 2 days. Finally, colonies from the source plates were transferred onto the assay agarose plates bearing different concentrations of metal salts using the ‘Replicate Many’ program of the Singer ROTOR HD^TM^ with the following settings: *recycle = Yes, revisit = Yes, source_pressure = 40%, source_pin_speed = 15 mm/s, source_overshoot = 1.5 mm, target_pressure = 25%, target_pin_speed = 13mm/s, target_overshoot = 1.2 mm, target_max = No, source_mix = No.* After 2 days of incubation at 30°C, all plates were scanned on an Epson V800 PHOTO scanner in grey and transmission scanning mode at 600 dpi using pyphe ^82^.

### QUANTIFICATION AND STATISTICAL ANALYSIS

#### Analysis of growth curves of WT cells

Data from the TECAN Spark-Stacker were processed in R. OD_600_ values of blank wells were subtracted from all sample wells before fitting sigmoidal growth curves using the *growthCurver* R package.

#### Analysis of ICP-MS data

Raw data from the Agilent 7700 ICP-MS were processed using Agilent MassHunter^TM^. Metal concentration (in parts per billion (PPB)) in each cultivation media as well as cell digest was calculated in MassHunter^TM^ using measurements of the calibrants (a 17-point dilution series of certified element standards) and scandium (Sc) as the internal standard to correct for minor deviations of the instrument during measurement. These values were processed further in R. For the cultivation media measurements, data from each media were compared with values in the control media and visualised. For the cell digests, after correcting for minor deviations in the total volume that dried pellets on the PVDF membrane from each batch was resuspended in (**Supplementary Table M5**), PPB values of each element in the blank samples was used to determine the limit of Quantification (LOQ). LOQ was defined as mean (PPB) + 5*sd(PPB) of the signal of each metal in the blanks of each individual batch. For all elements other than Sodium (Na) in batches 2-8 and Copper in batch 4, the quality control sample had a mean(PPB) > LOQ and in total 23 samples of a total of 342 were filtered out from the cellular metallomics data (**Supplementary Figure 1f**). To correct for varying cell numbers in the cell digests, we compared phosphorus (P) and OD_600_ normalisation strategies and discovered that P normalised data had fewer variation across biological replicates. Therefore, after filtering based on LOQ, the phosphorus (P) signal (which was observed to be stable and dependent on cell count) was used to normalise all other metals and the PPB values were scaled up to the original scale using the mean PPB values of the control samples. (Normalised PPB(m, x) = PPB(M,x)/PPB(P,x) * mean(PPB(P,x belongs to SM control). Batch correction was carried out such that the median value of each metal in the control samples was the same across batches. Data from samples with OD_600_ values > 0.1 were discarded. Nanograms per well values (1ng/ml = 1 PPB) after phosphorus normalisation were used to compare metal quantities across samples. For buffering capacity calculations, the measured metal concentrations in cultivation media were combined with those measured in cell digests and each set was normalised to the metal quantities in control samples (synthetic minimal media and cells cultivated in synthetic minimal media, respectively).

To estimate atoms per cell, cell number was estimated using OD_600_ values as described in ^83^. Atoms per cell = (6.022*10^23^ / atomic mass) *(pg per cell_BC_ * 10^-12^), where pg per cell_BC_ = (pg per cell / median (pg per cell in controls of batch) ) * median (pg per cell across all batches) and pg per cell = (PPB of metal (ng/mL) *1000) / (OD_600_ * 1.8 * 10^7^ * volume of culture transferred to each filter plate). The average coefficient of variation (CoV) across biological replicates of the ng/mL of digested cell cultures was 0.042, CoV in biological replicates of control samples was 0.032 (**Supplementary Table M6**) and the CoV of picogram per cell estimations across biological replicates of control samples was 0.13 (**Supplementary Table M7**), indicating again, that our OD_600_ estimates are likely more noisy than the ICP-MS measurements and better cell counting methods are required to obtain reliable estimations of atoms of each metal per cell. For most metals, the obtained cellular concentrations are consistent with previous studies. Only for Ca and Mn our concentrations values are slightly higher than in previous reports (**Supplementary Figure 1f**)

#### Processing of LC-MS data

All raw proteomics data (.wiff files) were processed using DIA-NN (Data-Independent Acquisition by Neural Networks ^41,84^) Version 1.8 compiled on 28 June 2021. The DIA spectral library (available at http://proteomecentral.proteomexchange.org/ , dataset ID PXD036062 (^24^ and resubmitted to proteomeXchange with dataset from this study) and FASTA file (UniProt yeast canonical proteome, downloadable from https://ftp.uniprot.org/pub/databases/uniprot/current_release/knowledgebase/pan_proteomes/UP 000002311.fasta.gz) used are identical to those generated for and described in ^24^. DIA-NN parameters used to process data are described in detail in **Supplementary Note M2**.

#### Normalisation, batch correction, filtering, and protein quantification

Quality control metrics exported by DIA-NN ( number of identified precursors > 0.4*max(number of precursors identified in any individual file), number of proteins identified >1000, total signal quantity > 1000000, MS1 signal quantity > 1000000, MS2 signal quantity > 10000000, normalisation instability < 0.5, Proteotypic ==1, Q value < 0.01, GG Q Value <= 0.01 and PG Q value < 0.01 ), optical density measured at the end of cultivation (OD_600_ units sampled > 0.75) and manual inspection of the total ion chromatograms of certain problematic files (26 files) were used to filter data processed by DIA-NN. In addition, 11 conditions (Cu 50, Cu 100, Fe 100, K 10, Mg 20, Mg 50, Mo 20, Mo 50, Zn 2e-04, Zn 0.001, Zn 0.002) were deemed as unsuitable for inclusion based on growth rate and metallomics measurements of media. Samples corresponding to perturbations of H_3_BO_4_ that were acquired and processed with the dataset were excluded at this stage to focus the study only on metals. In total, resulted in the retention of a total of 266 proteome data files out of the 437 that were acquired.

Protein quantities were estimated from peptide quantities using maxLFQ (using the DIA-NN R function diann_maxlfq()). Batch correction (median scaling) was carried out using median protein quantities of all control samples (WT yeast cells cultivated in synthetic minimal media). No imputation was carried out before the statistical analysis unless otherwise stated in the sections below. After all processing steps, the average replicate CoVs for perturbation condition samples was ∼15.7% with an average of 1837 proteins quantified per sample and for the control samples alone, the replicate CoV was ∼15.7% with an average of 1871 proteins quantified per sample.

Protein mass values were downloaded from the UniProt database (on 4th February 2024) and protein copy number values from ^42^ were combined with these to calculate what fraction of the protein mass of the 3841 proteins for which protein copy number data was available were measured or significant along any metal perturbation or cellular metal concentration series.

#### Identification of proteins differentially abundant along environmental metal concentration

Linear models with 0 (null model), 1, 2, and 3 degrees of freedom were fitted for each protein, modelling protein abundance as a function of measured metal concentration in cultivation media i.e. protein abundance ∼ poly (metal concentration , dof), where dof = 0, 1, 2 or 3. Only those protein - metal combinations which had at least 4 distinct points along the concentration gradient (rounded to 3 decimal spaces) and at least a total of 8 individual protein abundance measurements were used to fit the linear models. A series of F-tests were conducted using the anova() R function between all combinations of fitted models. P values from each F-test were adjusted using the Benjamini-Hochberg correction for multiple testing. To choose the simplest model that explains the data, the following logic was used: if the cubic (dof=3) model significantly outperformed all other models (dof=2, dof=1, and dof=0) i.e., adjusted p-value < 0.05, the cubic polynomial model was chosen. In cases where the cubic model did not outperform the linear model (dof=1), and the quadratic model (dof=2) did not surpass the linear model, but the linear model was better than the null model, the linear model was preferred. If the cubic model was not better than the linear model (dof=1), and the quadratic model (dof=2) was not better than the linear model, but the linear model was better than the null model, the linear model was selected. Next, if the cubic model was not better than the quadratic model, and the quadratic model was better than both the linear and null models, the quadratic model was chosen. Finally, if none of the cubic, quadratic and linear models performed better than the null model, the null model was selected as the simplest model. After selection of the least complex model, an additional threshold was applied for determining significant differential abundance along the metal perturbation series : proteins with a magnitude of fold difference (relative to control sample) change along metal perturbation series of at least 50% ( i.e. abs(max(fold difference along metal) - min(fold difference along metal)) > log_2_(1.5)) and P value of the simplest model to explain the expression pattern < 0.05 were deemed significantly affected.

#### Identification of proteins differentially abundant along cellular metal concentration

To identify protein differentially abundant along measure cellular metal concentration, relative metal quantification values from all data (for eg. Fe values from Fe perturbation as well as along Mg, Zn, Ca etc perturbations) were binned into bins of size 0.01 (metal concentration in each sample was normalised to that in control samples and rounded off to two decimal places). For each measured metal and each protein, the median of protein abundance values across the entire dataset corresponding to each bin along the measured cellular metal concentration was then modelled as a function of the measured cellular metal concentration using the same methodology as described above for identifying significantly differentially abundant proteins along environmental (media) metal concentration.

#### Correlation analysis using proteomics and metallomics profiles

Spearman’s rank-based correlation coefficient between each pair of samples within the metallomics dataset was computed in python 3 using the *scikit-learn* ^85^ library. The correlation coefficients were visualised as a heat map using the *seaborn* ^86^ library. The same methodology was followed for computing correlation coefficients between each pair of samples using proteomics data. The correlation coefficients computed based on proteomes and metallomes were then compared using *pearsonr* and *spearmanr* functions from *scipy.stats* ^87^.

#### Focused analysis of metal-related proteins

Gene function annotations in the gene ontology - molecular function (GO-MF) database (annotations fetched using the *AnnotationDbi* ^88,89^, *org.Sc.sgd.db* ^90^ and *GO.db* ^91^ R libraries) were used to annotate ORFs as “metal binding proteins” , “metal transport proteins” or “other metal related proteins”. Metallochaperones which are classified both as metal-binders as well as metal-transporters were considered “metal transport proteins” because their main function is to facilitate the incorporation of a metal into other metal-requiring proteins. Proteins for which a metal-specific annotation exists in the GO-MF were annotated with that metal, those annotated for more than one metal were included in metal binding or metal transport lists for both metals and those that bore “metal binding” or “metal transport” annotations without any specific annotations for a metal were labelled “orphan”.

#### Gene set enrichment analysis

All gene set enrichment analysis except for the network plots used to visualise clusters resulting from the ensemble clustering analysis in Figures 3d and 3e were performed in R using the *piano* ^92^ library and gene sets defined using the GO database (terms fetched as described above), KEGG database (fetched using the *KEGGREST*^93^ library) and GOslim annotations downloaded from the Saccharomyces Genome Database (http://sgd-archive.yeastgenome.org/curation/literature/go_slim_mapping.tab). The *piano::runGSAhyper()* function was used to carry out the enrichment analysis with a gene set size limits of 3 (lower limit) and 400 (upper limit), using all ORFs quantified as the background and Benjamini-Hochberg as the method for correcting P values of enrichment for multiple testing. Gene set terms with *P* value (adjusted) < 0.05 were considered significant. The results were visualised as Sankey plots (**Figure 2k**, **Supplementary Figure 2g**, **Figure 3c**, **Supplementary Figure 4b**) using the *plotly* ^94^ R library. Gene set enrichments and visualisation for **Figured 3a** and **3b** were conducted using the aPEAR R library ^95^.

#### Ensemble clustering analysis

An ensemble clustering framework was set up in *Python 3.9.13* (*numpy* 1.22.4 ^96^ , *scikit-learn* 1.1.1^85^, *igraph* 0.9.9 ^97^, *leidenalg* 0.8.9 ^54,97^, *seaborn 0.12.0* ^86^ and *scipy* 1.8.1 ^98^), in accordance with guidelines in ^49^ Briefly, the proteomics data were clustered in two parallel branches. The first, which we call *allmetal-clustering,* included proteins that were detected in at least 85% of all samples. The missing values in this dataset were imputed using the following imputation strategy: if measured quantities for a protein were missing in all samples in a metal perturbation condition (eg. Fe depletion (all samples cultivated in media containing lower Fe concentration than control (synthetic minimal) media), then the missing values for the protein in each sample corresponding to this perturbation were replaced with the minimum quantity of the protein detected in the entire dataset, if the protein was measured in at least one sample in the metal perturbation but missing in all replicates of a specific condition (eg. protein detected in Fe 0.5, but absent in all replicates of Fe 0.1), the missing values were replaced with the median quantity measured for the protein in all the control samples (cells cultivated in synthetic minimal media). Finally, if the quantity of a protein was missing in only some replicates of a cultivation condition (eg. two replicates of Fe 0.5 have missing values for a protein while the remaining two do not), the missing values were replaced by the median of the protein quantity of the replicates for which protein quantities were available.

For the second branch of ensemble clustering, called *metalwise-clustering,* a completeness filter of detection in at least 60% of all samples along each metal perturbation series was used before the imputation was carried out as described above. However, an additional filter was applied before performing the ensemble clustering analysis - only those proteins that were detected as being differentially abundant along either the environmental (media) metal concentration or along the measured cellular metal concentration were retained.

Three separate clustering algorithms were used to cluster the proteomics data within each parallel clustering pipeline:

● CommonNN ^50,51,99,100^ (the parameter R was varied equal-distant between 0.5\**R_cut* and *R_cut*, where R_cut was defined as the distance at which around 5% (2.5% for Mg and Mn due to smaller data set size) of the distances are smaller than *R_cut* (10 Rs), the number of shared nearest neighbours *N* was varied between 2 and 10 (step size 1) (9 Ns), the minimal cluster size *M* was set to 5. In total 90 R, N- combinations were used)
● kMeans++ ^52,53,85^ (the cluster number *k* was varied between 10 and 98 with a step size of 2 (between 10 and 55 with a step size of 1 for Mg and Mn). 45 clustering steps)
● Leiden ^54,97^ (the graph was set up using the 10 closest neighbours for each data point, with edge weights of 1-scaled(distance), all distances were scaled between 0 and 1).

For reproducibility, the seed was fixed to 42 for all clustering analyses. For each clustering algorithm a co-clustering matrix was calculated, where each element denotes the probability that two data points were clustered together. The co-clustering matrices from each clustering algorithm were combined into a single matrix, using equal weights. The final clusters were obtained by hierarchical clustering of the combined co-clustering matrix (Ward clustering ^86,98^), using a linkage-based cutoff for cluster extraction. Results from the clustering analysis were exported to R for gene-set enrichment analysis and visualisation as described above. Functions available at: https://github.com/OliverLemke/ensemble_clustering.

#### Analysis of growth screen of knock-out deletion mutants

Images of agarose plates from the Epson V800 PHOTO scanner were processed using the *gitter* R library to extract colony size. The .dat output from *gitter* ^101^ were combined with the experiment design table and analysed further ( grid normalisation, data aggregation, quality control checks and statistical analysis to obtain effect sizes and P values) using *pyphe* ^58,82^. Only 1% of the negative control positions (footprints) (49 out of 4875 empty spots in total across all agarose plates) were contaminated and no systematic contamination was observed. One plate (corresponding to the reduction of potassium to 1/50^th^ the level in synthetic minimal agarose media) contained 13 contaminated footprints and was therefore excluded from further analysis. Correlation between replicates of the control strain within a single plate was 0.78 before for the raw colony sizes and 0.95 after correction for surface effects using the control strain grid. Next, *pyphe-interpret* to obtain effect sizes of each mutant relative to the control strain (*Δhis3* from the haploid prototrophic library we used) and *P* values from Welch’s t-test for samples with unequal variance corrected for multiple testing using the Benjamini-Hochberg method. In total, 357972 colony size measurements, corresponding to 4759 unique deletion mutant (and control) strains across 17 cultivation media conditions, remained after data processing. A *P* value (adjusted) threshold of < 0.10 and an abs(log_2_(effect size)) > log_2_(1.2) was chosen to determine which mutants showed a significantly altered growth in each condition. Data from different levels of depletions of K, Mg and Zn and from Fe depletion and Fe depletion combined with the dipyridyl chelator were combined at this stage to determine the final list of ORF deletions that were affected to enable a metal-wise comparison with all other datasets.

#### Defining metal-related genes

The Gene Ontology database ^91^, specifically the Molecular Function GO annotations were used to determine a set of metal-binding proteins, metal-transport proteins and other metal-related genes that do not fall under the first two categories (eg. “calcium-dependent protein kinase C activity”). Open reading frames (ORFs) mapping to GO terms containing the word “binding” and any word reflecting involvement of Ca, Cu, Fe, Mg, Mn, Mo, Na, Zn, heme or protoheme were included in the metal-binding annotation set with specific annotations. ORFs mapping to terms containing the words “metal binding” but no indication of which metal or metal-containing group is bound were annotated as “orphan”.

Metal transporters and other metal-related ORFs were filtered from the GO-MF database using a manual creation process and text parsing using regular expressions was not sufficient to include only metal related transporters or the metal dependent enzyme activities included in the “other metal related” set. A list of these terms is available at *input_processed_databases_publisheddatasets.R* within the code repository at https://github.com/Ralser-lab/metallica.

#### Comparison and integration with published datasets

We did not modify metallomics data from mutant strains before using and directly used Z-scores of cellular metal concentration in each deletion and overexpression mutants reported by Iacovacci et al. ^27^, which were calculated using metal concentration measurements collected by Danku et al ^26^ (cellular metal concentration in each haploid knockout mutant of *S. cerevisiae)* and Yu et al. ^25^ (overexpression mutants). We annotated any Z-score with an absolute value > 1.959 (corresponding to p-value < 0.05) as a significant change in metal concentration in a mutant.

Protein abundance data in haploid knockout deletion mutants were sourced from Messner et al ^24^. We filtered this dataset to retain only knockouts of genes known to be connected to specific metals in the GO-MF database-based mapping described above and considered any protein quantified in these mutants with absolute log_2_(fold difference vs. control) > log_2_(1.5) and p-value < 0.05 as being significantly altered. The upset plot (**Figure 4h**) to visualise commonalities between genes identified as connected to metals were created using the upSet R package. UniProt annotation status annotations were merged with all the datasets to assess how many poorly characterised genes were identified as significantly affected in each dataset (**Figure 5a-c**). The circular plot to summarise current annotation status, metal-binding and metal transporter annotations and all the datasets (**Figure 5c**) was created using the *circos* R library. Gene set enrichments for Supplementary Figure 4b were conducted and visualised as described above.

#### Simulations of metabolic flux using CofactorYeast

Simulations of excess and depletion of each metal were carried out in MATLAB using the *cobratoolbox* ^102,103^ and the CofactorYeast framework ^66^. CofactorYeast incorporates metal ion cofactors and import and export reactions for each metal ion into the Yeast8 genome-scale metabolic model ^104^. Growth rates were fixed to the lowest experimentally measured growth rate upon each metal perturbation (eg. Fe depletion or Zn excess). Protein abundances of each enzyme were allowed to vary to achieve the minimisation or maximisation of the metal uptake. Flux balance analysis was carried out to simulate the fluxes required to achieve the objective function (minimisation or maximisation of the metal transport) under the growth rate constraint. The flux results (**Supplementary Information - Results**) were then processed in R to calculate flux change values (flux in perturbation condition / flux in control condition). This resulted in several infinite values due to flux = 0 of either the control condition simulation or the perturbation condition simulation. Therefore, infinite flux changes values of conditions for which both the control condition and simulation condition flux was 0, was set to 1. For conditions where the control condition flux was not zero, but the simulated flux was zero were set to a low value with the correct direction (i.e., *log_2_(fold change flux) = -sign(control condition flux)* and for those where the control condition flux was zero, but the perturbation flux was nonzero, it was set to *log_2_(fold change flux) = sign(perturbation flux)*. Reactions through which abs(log_2_(fold change flux)) > log_2_(1.5) were considered significantly affected.

#### Metabolic network analysis using *igraph*

The Yeast8 metabolic model was downloaded from https://github.com/SysBioChalmers/yeast-GEM and used as the input to create a directed, bipartite graph using the *igraph* python library. Reaction IDs and metabolite IDs were used as the two types of nodes. Directed edges between the nodes were from each substrate metabolite node to each reaction node and from each reaction node to each product node. Since 1670 of the 4131 reactions in the Yeast8 model were reversible, directed edges were added to the graph in both directions for these. We noticed a slight imbalance in the fraction of reversible reaction nodes mapping to at least one metal-linked enzyme and those that did not: only 21.38% of the metal-linked reaction nodes were reversible while 43.71% of those without metal-linked annotations were reversible. Therefore, all our calculations on the *igraph* that are grouped using the metal-linked annotation have a slight bias of counting nodes without metal annotations more often than metal-requiring nodes. The final graph was not a fully connected graph. However, because 99.55% of nodes would be retained if we filtered the graph for its largest weakly connected component and 88.47% nodes would be retained in the largest strongly connected component, we proceeded without filtering the graph.

Weak or strongly connected components were determined by the *clusters (mode = “weak”)* and *clusters(mode = “strong”)* functions from the *igraph* python library. The graph was visualised, and all centrality metrics were calculated using *igraph* functions.

## Data visualisation

All data visualisations except **Figure 6b**, **6c** and **6g** were created in R using the ggplot2 ^105^, plotly ^94^, viridis^106^, aPEAR, RColorBrewer ^107^ and circlize^108^. The bipartite, directed network in **Figure 6c** was visualised using the *igraph* ^109^ python library and **Figures 6b** and **6g** were created using iPATH ^110^. Adobe illustrator and Biorender were used to assemble some figures.

## Lead contact

Further information and requests for resources and reagents should be directed to and will be fulfilled by the lead contact, Markus Ralser.

## Materials availability

Requests for reagents should be directed to and will be fulfilled by the lead contact.

## Data and code availability

All supplementary information and result data files are available at Zenodo (DOI:**10.5281/zenodo.10708992**). Raw data and code used to analyse data will be available after peer review.

## Acknowledgements

We thank Benjamin Heineike, Christoph Messner, Lucia Herrera-Dominguez, Clara Correia-Melo, Enrica Calvani for support throughout the execution of this project, James Macrae, Luiz Carvalho and Jürg Bähler for key input, and Gavin Kelly from the Bioinformatics and Biostatistics STP at the Francis Crick Institute for key input for the statistical analysis of proteomics data. This work was supported by the Francis Crick Institute, which receives its core funding from Cancer Research UK (no. FC001134), the UK Medical Research Council (no. FC001134), the Wellcome Trust (no. FC001134 and IA 200829/Z/16/Z), and the European Research council as part of the ERC-SyG-2020 (951475).

## Author Contributions

S.K.A and M.R. conceptualised and designed the study. S.K.A. and J.H. conducted growth rate measurements. S.K.A and L.S. collected and analysed ICP-MS measurements. S.K. & S.K.A. designed, conducted and analysed the knockout deletion mutant growth screen. O.L. performed ensemble clustering analysis. Y. C. simulated metal perturbations using flux balance analysis.

S.K.A performed all other data analysis and visualisation. M.M. provided key input for designing the study. S.K.A. and M.R. wrote the manuscript. All authors reviewed and revised the manuscript.

## Supplementary Figures

**Supplementary Figure 1.**
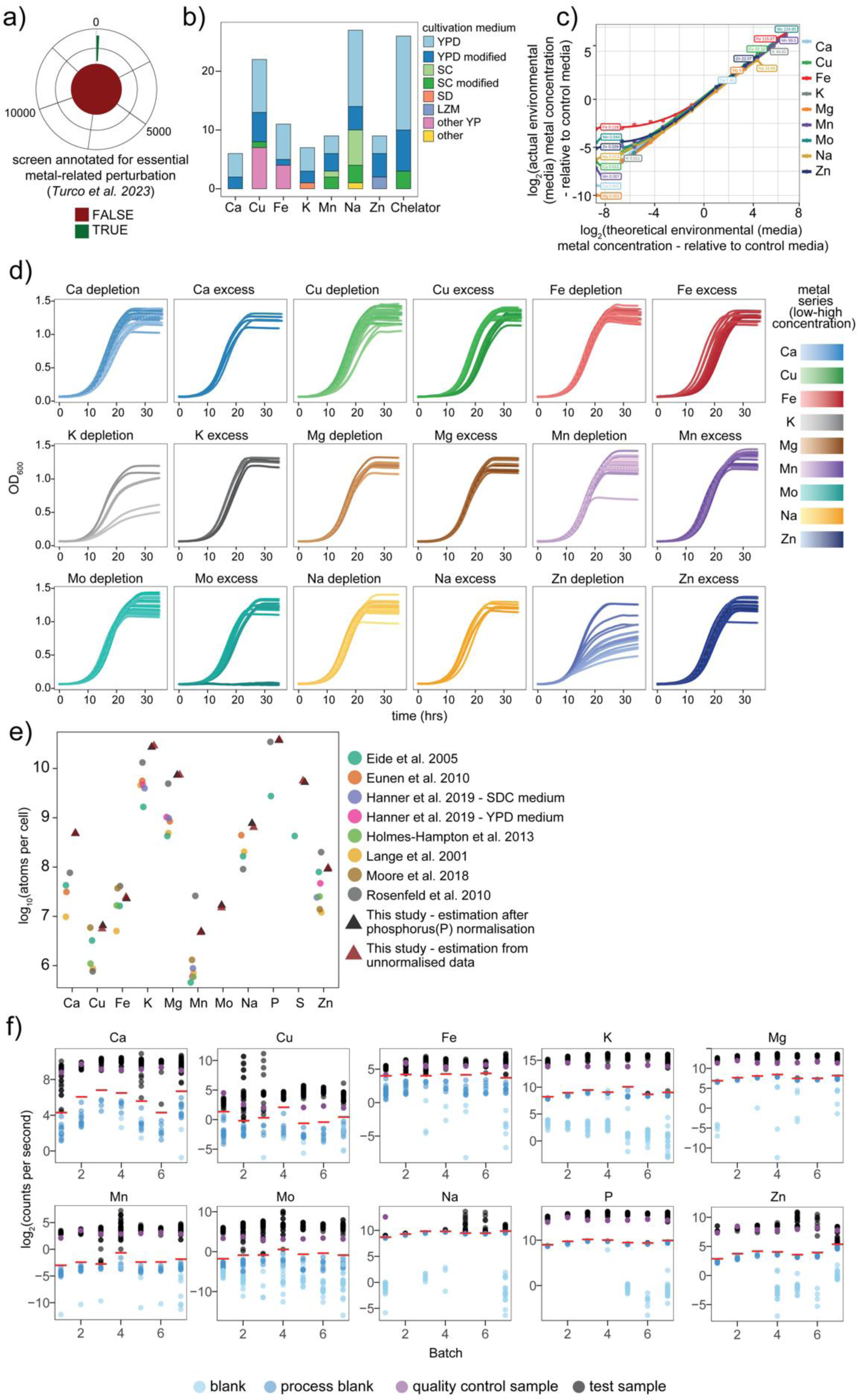
a) The number of experiments compiled by ^12^ that represent concentration perturbations of physiological relevant metal ions (green) compared to all other types of environmental perturbations (red). b) Most experiments corresponding to concentration perturbations of physiologically relevant metals were conducted in rich media. X-axis: physiologically relevant metal or chelator condition, Y-axis: number of experiments compiled by ^12^ in which the metal was perturbed. Colour indicates the type of media that was used as the base for perturbations (YPD: Yeast extract peptone dextrose, SC: synthetic complete, SD: synthetic defined (same as synthetic minimal media), LZM: Low zinc media, other YP: media composed of yeast extract and peptones but a different sugar source). c) Concentration of metal ions in each metal perturbation media as quantified by Inductively Coupled Plasma - Mass Spectrometry (ICP-MS). X- axis: log_2_(theoretical metal concentration in each cultivation media - relative to synthetic minimal (control) media). Y-axis: log_2_(ICP-MS based measured concentration of each cultivation media - relative to synthetic minimal (control) media). Colour indicates metal being perturbed in media. Labels indicate the lowest and the highest relative metal concentrations that were measured. d) Growth curves of *Saccharomyces cerevisiae* (WT BY4741+pHLUM) cells in media with perturbed metal concentration. X-axis: time (in hours), Y-axis: Optical Density (OD_600_). Colour indicates the media. Intensity of colour indicate the concentration of metal (light - dark corresponding to low - high concentrations) e) Comparison of estimation of total cellular metal concentration of each metal quantified in this study and previous reports. X-axis: metal. Y-axis: log_2_(ICP-MS or ICP-AES based estimation of atoms per cell). Colour indicates the study. f) ICP-MS data collected across all batches for blanks (light blue), process blanks (darker blue), quality control samples (purple) and test samples (black).X-axis: batch number, Y-axis: log_2_(counts per second as measured by ICP-MS). Red lines indicate LOQ (limit of quantification) = mean(blanks) +- 10*sd(blanks).

**Supplementary Figure 2.**
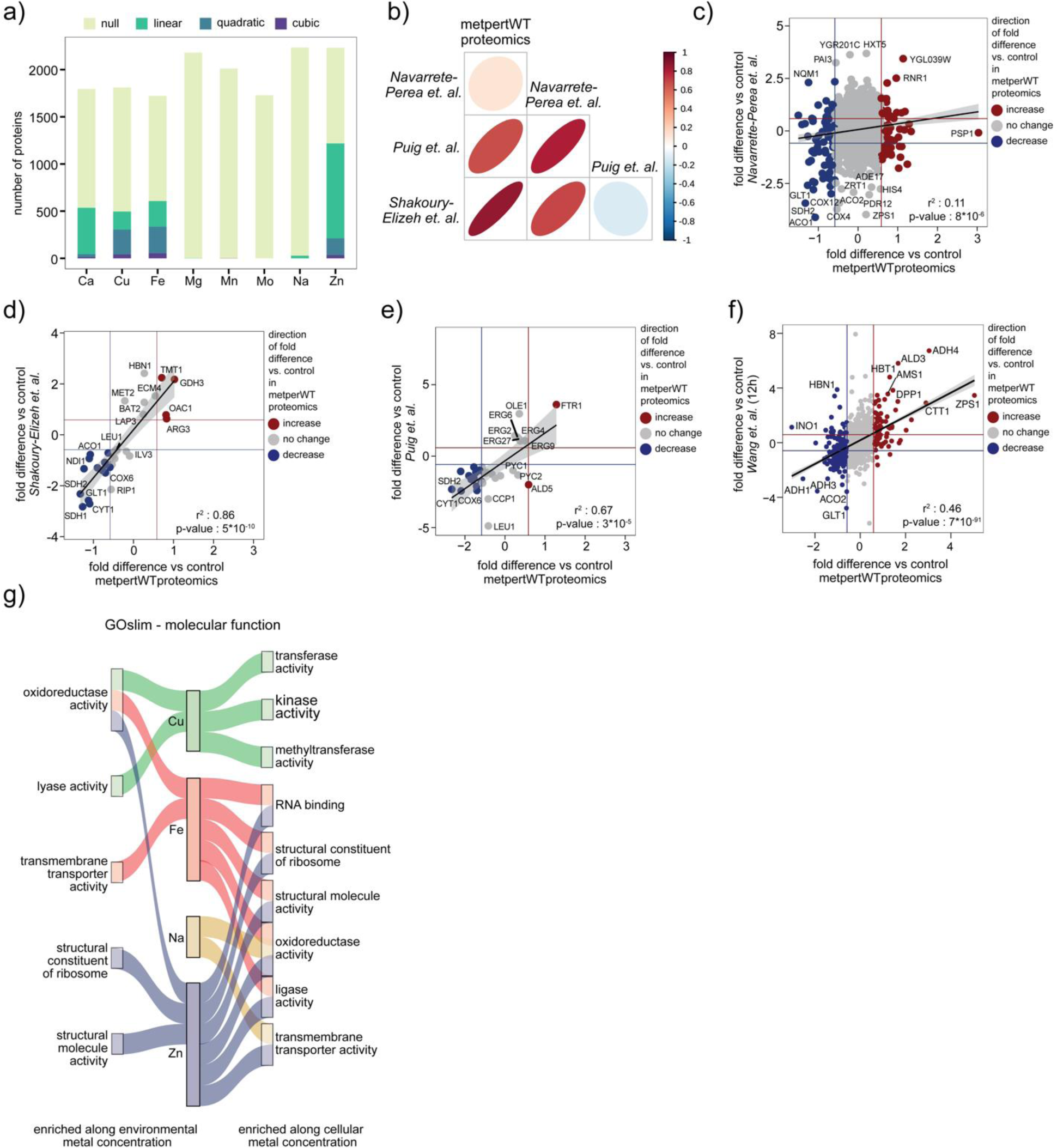
a) Number of proteins classified based as having no significant abundance change (null model) or following a linear, quadratic or cubic protein abundance profile along environmental concentration of each metal based on the statistical analysis (F-test p-value < 0.05). X-axis: metal being perturbed in the environment. Y-axis: number of proteins. Colour indicates the type of linear model that was deemed as the simplest model that explains the relationship between protein abundance and environmental metal concentration. |b) Summary of correlation between the quantitative proteomics dataset described in this study and previous work on Fe depletion. Right leaning ellipses and red colour indicate a positive correlation. Left leaning ellipses and blue colour indicate a negative correlation. c) - e) Correlation between quantitative transcriptome or protein abundance data from Fe depletion samples acquired in this study and three others, namely, ^16^ (c) , ^15^ d) and ^13^ (e). f) Correlation between quantitative transcriptome or protein abundance data from Zn depletion samples acquired in this study and ^17^). g) GOslim - molecular function terms enriched in groups of proteins significantly differentially abundant along each environmental (left) and cellular (right) metal perturbation series. Colour indicates metal that was perturbed in the environment or measured in cells.

**Supplementary Figure 3.**
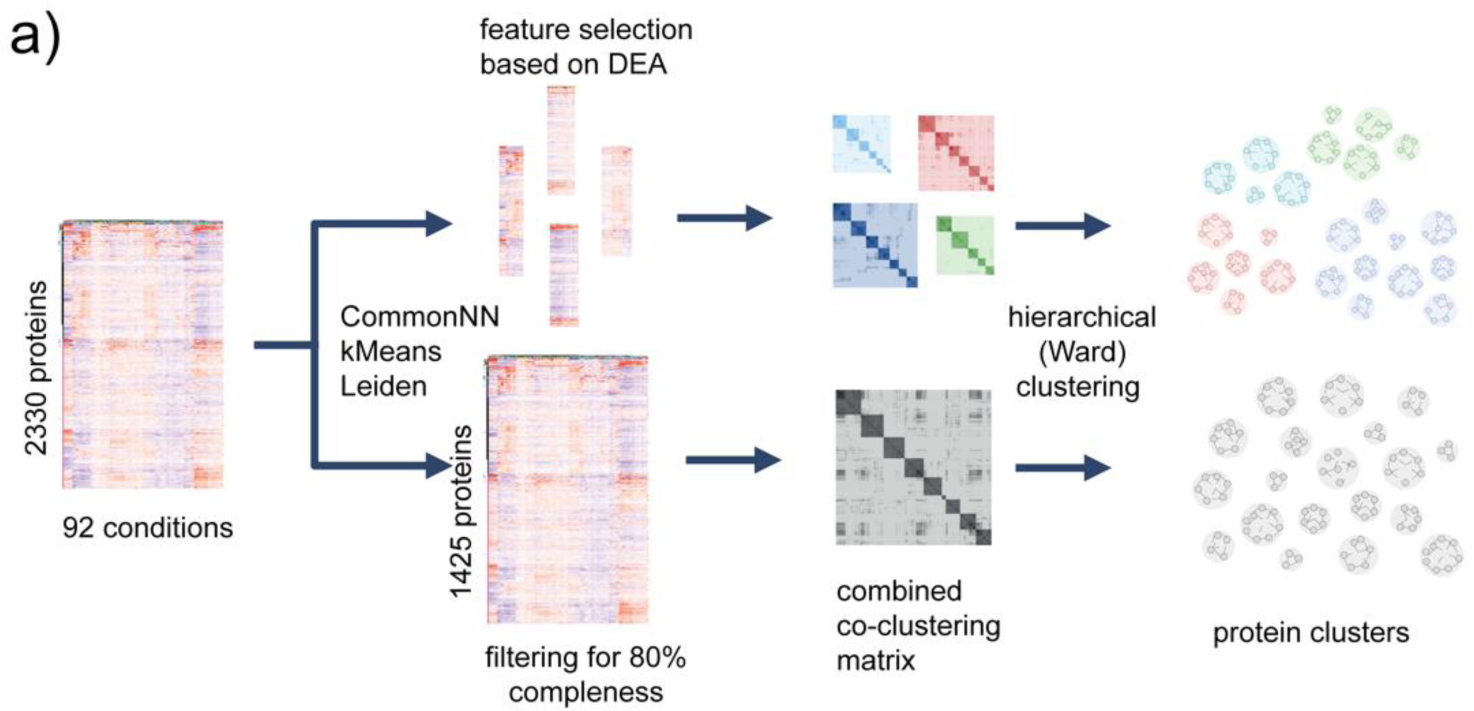
a) Visual summary of the ensemble clustering pipeline.

**Supplementary Figure 4.**
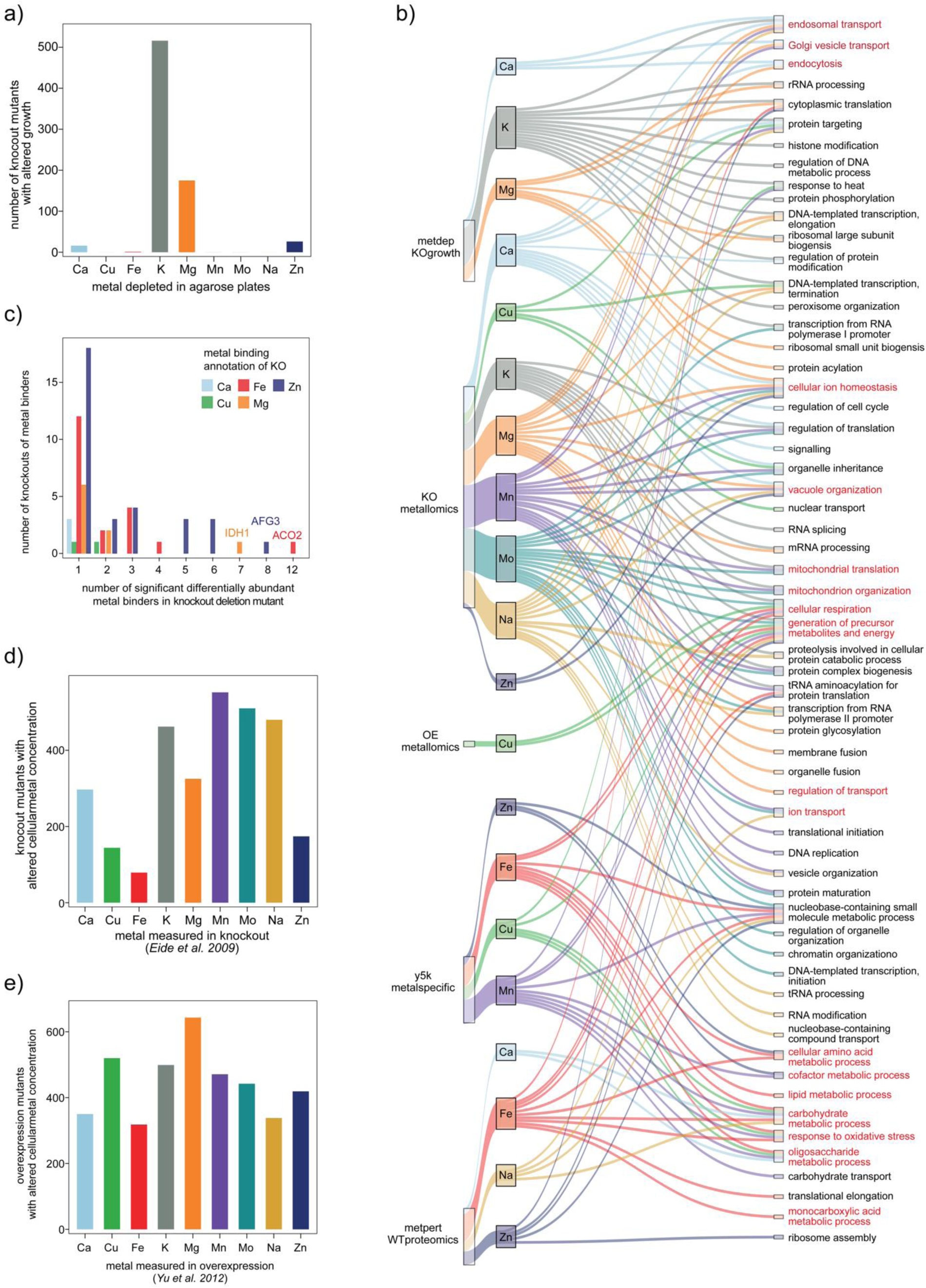
a) Number of metal-gene interactions identified by cultivating the prototrophic haploid knockout (BY4741+pHLUM) collection of *S. cerevisiae* on agarose plates bearing depleted amounts of metals. X-axis: metal that was depleted in agarose plates. Y-axis: Number of genetic interactions identified (based on reduction or increase in growth of each mutant) b) GOslim - biological process terms enriched in genetic interactions with metals or differentially abundant proteins identified in each dataset. Colours indicate the metal for which a process was enriched in the interacting genes or differentially abundant proteins. c) Number of metal-gene interactions identified based on metallomics data from S. cerevisiae knockout mutants acquired by Eide et al 2009 and analysed by Iacovacci et al 2021. X-axis: metal that was quantified in each mutant. Y-axis: Number of genetic interactions identified. d) Number of metal-gene interactions identified based on metallomics data from S. cerevisiae overexpression mutants acquired by Yu et al 2012 and analysed by Iacovacci et al 2021. X-axis: metal that was quantified in each mutant. Y-axis: Number of genetic interactions identified. e) Distribution of metal binding proteins differentially abundant in knockout mutants of metal binding proteins. X-axis: number of differentially abundant proteins in a knockout. Y-axis: number of knockouts. Colour indicates the metal binding annotation of the protein encoded by the knocked-out gene.

**Supplementary Figure 5.**
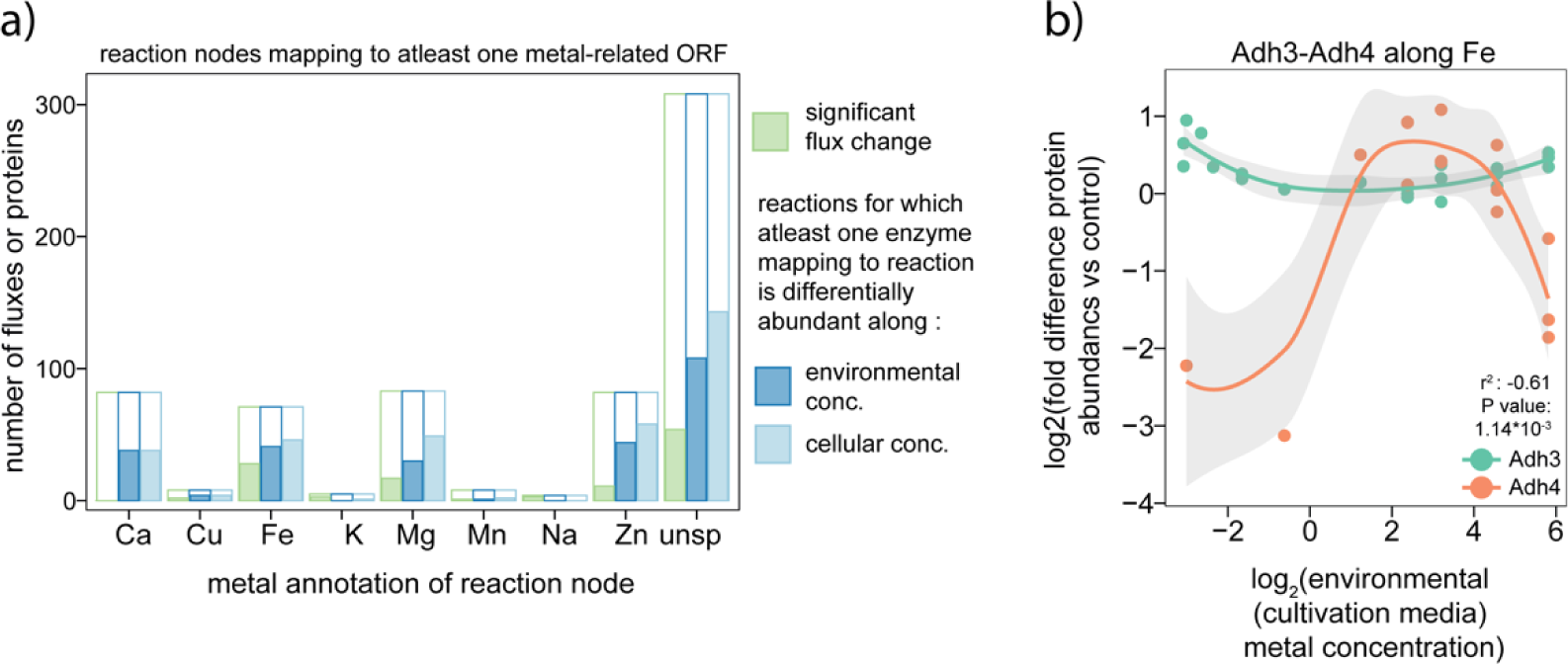
a) Significantly altered fluxes based on FBA simulations (green), differentially abundant proteins identified along environmental metal concentration (blue) and cellular metal concentration (light blue) at reaction nodes bearing at least one metal binding annotation. X-axis: metal that is annotated to bind to at least one enzyme that catalyses the reaction (“unsp”: unspecific - unclear which metal binds the enzyme). Y-axis: number of fluxes or proteins that were quantified (outer bar) and identified as significantly altered (filled up fraction of bar). b) Protein abundance profiles of the isozymes Adh3 (green, Zn- binding annotation only) and Adh4 (orange, Zn and Fe binding annotations) along environmental Fe concentration. X-axis: log_2_(environmental Fe concentration). Y-axis: log_2_(fold difference protein abundance vs control condition).

